# Two Competing Guilds as a Core Microbiome Signature for Health Recovery

**DOI:** 10.1101/2022.05.02.490290

**Authors:** Guojun Wu, Ting Xu, Naisi Zhao, Yan Y. Lam, Xiaoying Ding, Dongqin Wei, Jian Fan, Yajuan Shi, Xiaofeng Li, Mi Li, Shenjie Ji, Xuejiao Wang, Huaqing Fu, Feng Zhang, Yu Shi, Chenhong Zhang, Yongde Peng, Liping Zhao

## Abstract

Over eons of co-evolution, the gut microbiota has become an essential organ for humans^1,2^. However, it is unclear what core members and their ecological organization ensures the stable provision of this organ’s essential health-relevant functions to the host. With high quality metagenome-assembled genomes as network nodes, here we identified two competing guilds^3^ of the most stably and highly connected bacteria that together correlate with a wide range of host health conditions. Genomes in these two guilds kept their ecological relationship unchanged despite experiencing profound abundance changes during a 3-month high fiber intervention and 1-year follow-up in patients with type 2 diabetes (T2DM). The genomes of one guild harbored more genes for plant polysaccharide degradation and butyrate production, while the other guild had more genes for virulence or antibiotic resistance. A Random Forest regression model showed that the abundance distributions of these genomes were associated with 41 out of 43 bio-clinical parameters in the study cohort. With these genomes as reference, Random Forest modeling successfully classified case and control of T2DM, atherosclerotic cardiovascular disease, liver cirrhosis, inflammatory bowel diseases, colorectal cancer, ankylosing spondylitis, schizophrenia, and Parkinson’s disease in 12 independent metagenomic datasets from 1,816 participants across ethnicity and geography. This core microbiome signature may serve as a common target for health recovery.

## Introduction

The gut microbiome supports the host’s homeostasis in metabolism, immunity, development, and behavior, etc.^4^ It has been regarded as an essential organ because the attenuation or loss of health-relevant functions of a dysbiotic gut microbiome has been linked with the initiation and/or progression of many chronic diseases, including type 2 diabetes (T2DM)^5–7^. However, the core members of the gut microbiota and their underlying ecological structure remain to be identified.

The gut microbiota is a complex adaptive system^8^, in which the minimum responding units to environmental perturbations are bacterial genomes^9^. More importantly, genomes are not independent microbiome features. They form ecological interactions, such as competition or cooperation, with each other and organize themselves into a higher-level structure called “guilds”^3^. Each guild is a potential functional group of bacteria in the gut ecosystem. Guild members may have widely diverse taxonomic backgrounds but thrive or decline together and thus show co-abundant behavior. Guild-level variations have been positively or negatively correlated with disease phenotypes and their members have been demonstrated as having causal role in host disease phenotypes^10,11^. Although a suite of microbiome-wide association studies (MWAS) has attempted to identify the microbiome signatures (using features such as genes, pathways, taxa, etc.) that are associated with disease phenotypes ^12–15^, genomes and their guild-level organization have not been extensively employed to describe the ecological structure that supports the gut microbiota to stably provide health-relevant functions to the host. To this end, we adopt a genome-centric approach which is based on high-quality draft genomes assembled directly from metagenomic datasets (high-quality metagenome-assembled genomes, HQMAGs). This approach uses genomes as nodes of ecological networks and their guild-level aggregations as ecologically meaningful features for identifying microbiome signatures associated with host phenotypes in health and diseases.

In this study, we hypothesized that bacteria required for providing essential health-relevant functions to the host^2^ should maintain stable ecological interactions with each other to form robust guilds^16,17^. To identify microbiome signatures that are based on stable interactions among HQMAGs, we randomized T2DM patients at baseline (M0) to receive either 3 months (M3) of a high fiber intervention^11^ (W group; n = 74) or standard care (U group; n= 36) followed by a one-year follow-up (M15) in an open label, randomized, controlled trial (Fig. 1A and Fig. S1). The high fiber intervention was used to exert an environmental perturbation to dramatically change the abundance of members of the gut microbiome^10,11^. Co-abundance network analysis at each of the three time points enabled us to identify genome pairs that can keep their correlations unchanged despite significant community-wide abundance changes caused by the perturbations. We found that these robust genome pairs were from 141 HQMAGs and these genomes formed two guilds. These two guilds were organized as the two competing ends of a robust seesaw-like network, whenever one guild increased, the other decreased in abundance. Together, these seesaw networked genomes supported Random Forest models for predicting the response of a wide range of metabolic phenotypes to dietary intervention in the T2DM cohort, as well as for classifications of case and control of 8 different diseases in 12 independent metagenomic datasets suggesting that we may have identified a core microbiome signature across different chronic diseases.

**Fig.1.**
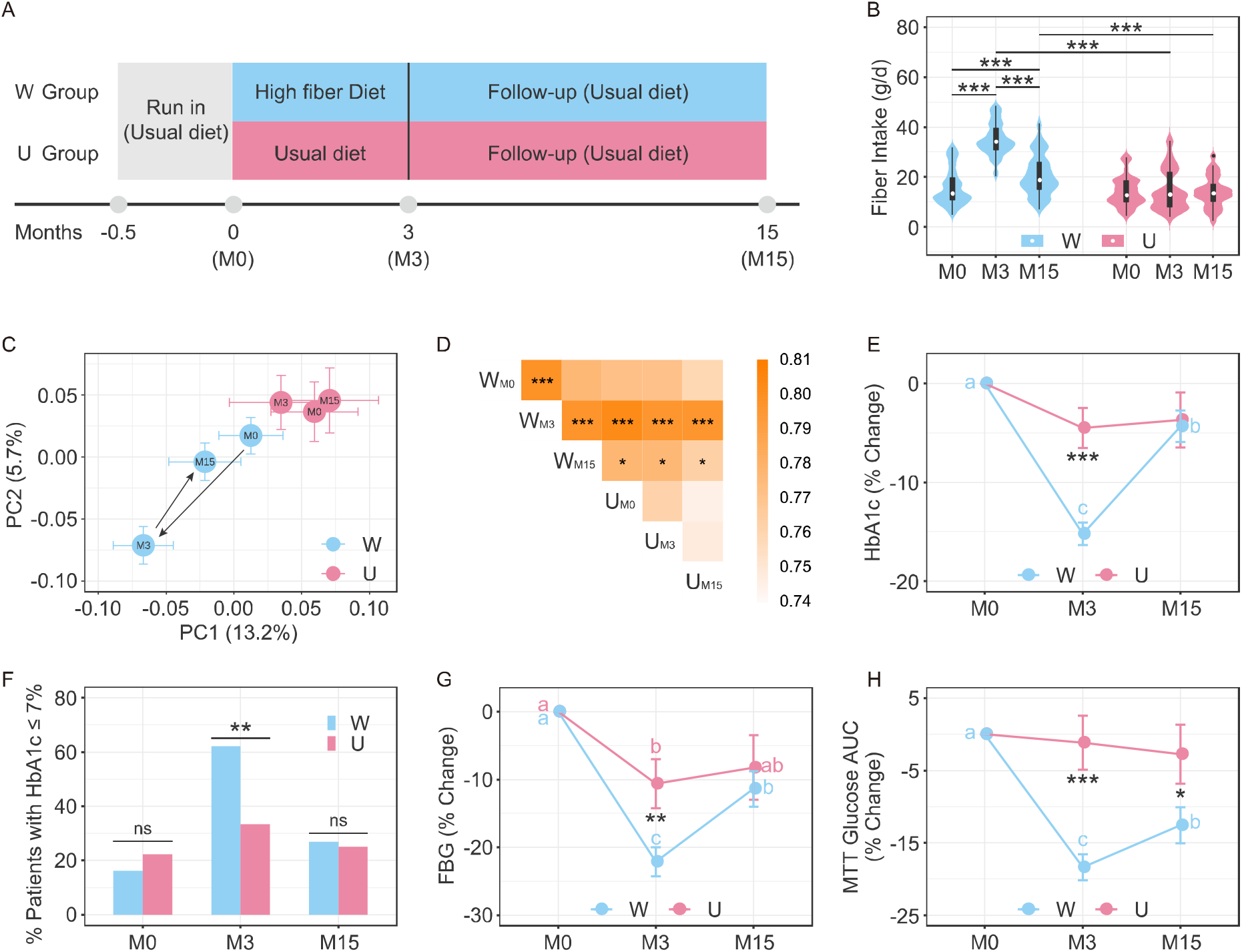
Reversible changes of gut microbiota associates with reversible shifts of metabolic phenotypes in patients with T2DM. (A) Study design. Before Run-in, written informed consent, questionnaire of personal information and measuring HbA1c at screening. After Run-in, medical checkup and sample collection at baseline (M0), three months after on the high fiber intervention or usual diet (M3) and one year after the high fiber intervention stopped (M15). (B) Changes of fiber intake. (C) Global changes of the gut microbiome as shown by the principal coordinate analysis based on the Bray-Curtis distance for the 1845 genomes and (D) Average Bray-Curtis distance between the groups. PERMANOVA test (9,999 permutations) was performed to compare the groups. * P < 0.05 and *** P < 0.001. The color of the square showed the magnitude of average Bray-Curtis distance. (E) Change of HbA1c, (F) The percentage of participants with adequate glycemic control, (G) Fasting blood glucose, and (H) The glucose area under the curve (AUC) in a meal tolerance test (MTT). For (E), (G) and (H), data shown as percent changes from baseline (± S.E.M). Friedman test followed by Nemenyi post-hoc test was used for comparison in the same group, compact letters reflect significance (P < 0.05). n = 67 in W group and n = 28 in U group. Mann-Whitney test (two-sided) was used for comparison between W and U at the same time point, * P < 0.05, ** P < 0.01 and *** P < 0.001. n = 74 in W (M0) (For panel H, n=72), n = 74 in W (M3), n = 67 in W (M15), n = 36 in U (M0), n = 36 in U (M3) and n = 28 in U (M15).

## Results

### Reversible changes in the gut microbiota associate with reversible changes of host metabolic phenotypes

Dietary fiber intake in the U group remained unchanged throughout the study, whereas W group had a significant increase in the intake of dietary fibers from M0 to M3 and a decrease from M3 to M15 (Fig. 1B). Compared with the U group, fiber intake was significantly higher in the W group at both M3 and M15 (Fig. 1B), but energy and macronutrient consumption were similar between the two groups across the study period (Fig. S2).

To investigate the gut microbial responses to the introduction and withdrawal of the high fiber intervention, we performed shotgun metagenomic sequencing on 315 fecal samples collected from 110 patients of the W and U group, among whom 95 patients provided samples at all 3 time points and 15 provided samples at M0 and M3 only (Table S2, Fig S1). To achieve genome-level resolution, we reconstructed 1,845 non-redundant high-quality draft genomes (HQMAGs, two HQMAGs were collapsed into one if the average nucleotide identity, ANI, between them was > 99%) from the metagenomic datasets. These HQMAGs accounted for more than 70% of the total reads. In the context of beta-diversity based on Bray-Curtis distance, the overall structure of the gut microbiota in the W group significantly changed from M0 to M3 (PERMANOVA test, P < 0.001) and returned to that of M0 at M15; there was no difference in the U group across the 3 timepoints (Fig. 1C, D). Similar changes in alpha-diversity based on Shannon and Simpson indices were also observed (Fig. S3). These results showed that the high fiber intervention induced significant structural changes of the gut microbiota^11^, however the gut microbiota reverted to baseline after the intervention was withdrawn indicating a high resilience in community structure.

To determine if host metabolic phenotypes would show similar reversible changes as the gut microbiota, we examined 43 bio-clinical parameters of 9 categories across the 3 time points (Table S3). Hemoglobin A1c (HbA1c) in the U group showed no significant changes throughout the trial. The high fiber intervention significantly reduced the level of HbA1c in the W group from M0 to M3 by 15.22 ± 9.82% (mean ± SD). At one-year follow-up of the W group, HbA1c was significantly increased from M3 but remained lower than that at M0 (Fig. 1E). The proportion of patients who achieved adequate glycemic control (HbA1c ≤ 7%) was significantly higher in the W group (61.6 % versus 33.3% in the U group) at M3, but showed no difference between the two groups at M15 (Fig. 1F). The level of fasting blood glucose and postprandial glucose in meal tolerance test followed a similar trend as HbA1c (Fig. 1G, H). Among the rest 40 bio-clinical parameters, 14 also showed an alleviation from M0 to M3 but rebounded at one-year follow-up in the W group (Table S3). These results indicate that changes of the host metabolic phenotypes were associated with the reversible changes of the gut microbiota in response to the introduction or withdrawal of the high fiber intervention.

### Genome pairs with stable interactions form a seesaw-like network of two competing guilds

To facilitate the identification of genome pairs that keep their ecological interactions stable during the trial, particularly in the W group with profound microbiota and host phenotypic changes, we constructed a co-abundance network for each time point based on the abundance matrix of the HQMAGs representing the prevalent microbes. Co-abundance network is a data-driven way to investigate ecological interactions between microbes across habitats^18,19^. A total of 477 HQMAGs were selected for network construction because they were detectable in more than 75% of the samples at each time point in the W group. These 477 HQMAGs also accounted for ~60% of the total abundance of the 1,845 HQMAGs. In the W group, we calculated pairwise correlations of all 113,526 possible genome pairs among these 477 prevalent HQMAGs based on their abundance across the patients at each time point and constructed 3 co-abundance networks (***G***_M0_, ***G***_M3_ and ***G***_M15_) (Figure 2A, Table S4). The three networks were of similar order ***S***, i.e., the total number of nodes (HQMAGs), ***S***_M0_(442), ***S***_M3_(421), and ***S***_M15_(429), but they varied considerably in their size ***L***, i.e., the total number of edges (correlations), ***L***_M0_(4,231), ***L***_M3_(2,587) and ***L***_M15_(4,592). ***L*** in ***G***_M3_ decreased to 61.14% of that in ***G***_M0_ and rebounded back in ***G***_M15_ to 108.53% of that in ***G***_M0_. This pattern was confirmed by changes in connectance, which is defined as the proportion of realized ecological interactions among the potential ones (in undirected network, connectance= 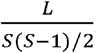, range: [0,1])^20^. Connectance decreased from 0.043 in ***G***_M0_ to 0.029 in ***G***_M3_ and rebounded to 0.050 in ***G***_M15_. Changes in L and connectance showed that the high fiber intervention dramatically reduced the correlations among the prevalent genomes in the network. In addition, we found that the distributions of degree, i.e. the number of edges a node has, fit well with a power-law model (Fig. S4, R2 values ***G***_M0_: 0.79, ***G***_M3_: 0.82, ***G***_M15_: 0.79), indicating the presence of network hubs^21^. Defining hubs as nodes that connect with more than one-fifth of the total nodes in the network (Fig. S5), we found 24 hubs, in which 10 were in ***G***_M0,_ 20 were in ***G***_M15_ but none were in ***G***_M3_. These results indicate that the overall structure of the gut microbiome undergone profound changes during the trial, particularly, the high fiber intervention resulted in the loss of interactions between genome pairs.

**Fig. 2.**
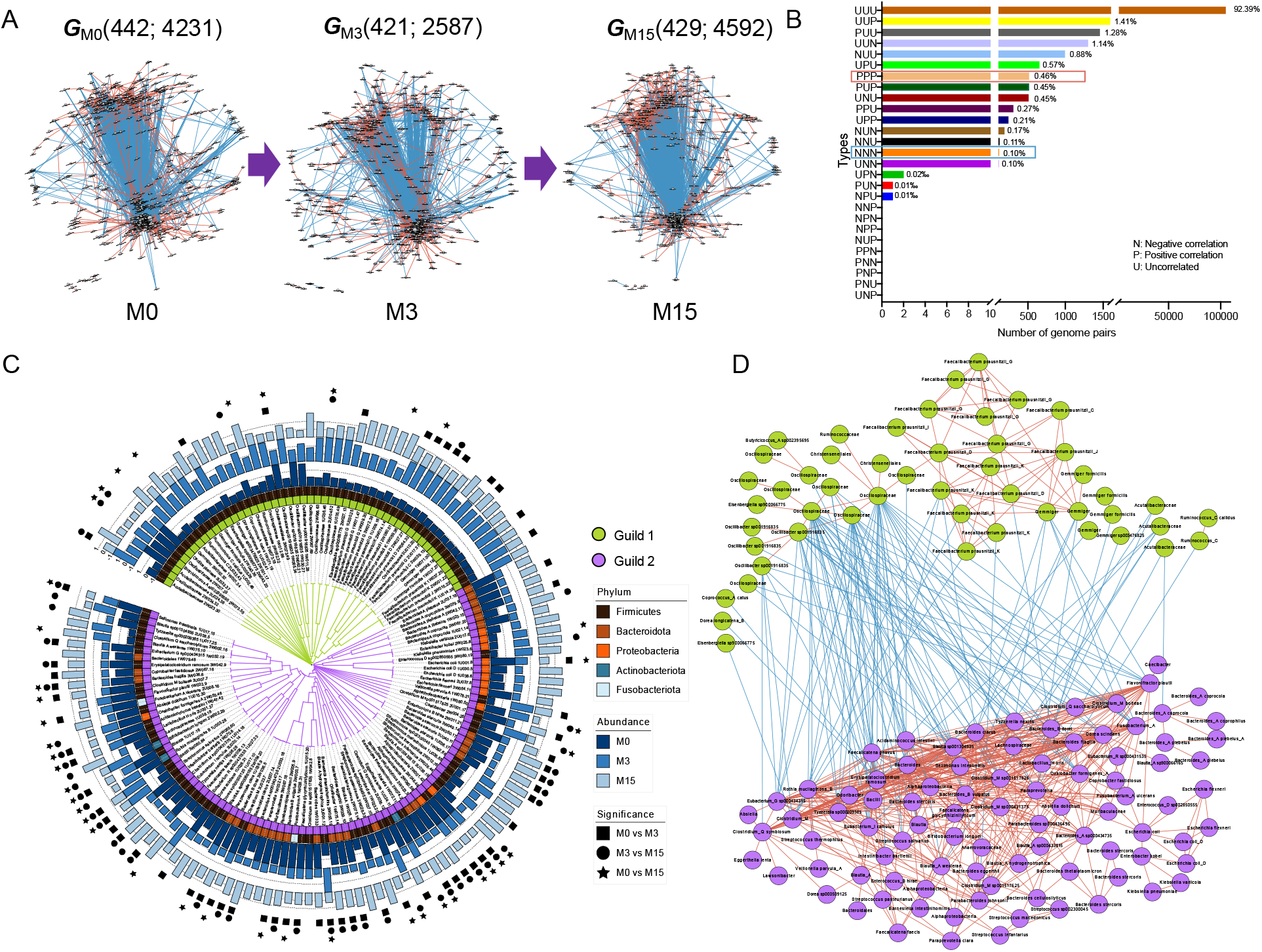
Two competing guilds of bacteria constitute a robust seesaw-like network despite the profound global changes in the gut microbial ecosystem induced by introduction and withdrawal of the high fiber intervention. (A) Co-abundance networks of the prevalent genomes in W group at M0, M3 and M15 during the trial, denoted as ***G***_M0_(442; 4231), ***G***_M3_(421; 2587) and ***G***_M15_(429; 4592), numbers in parenthesis are order and size of the network. The correlations between genomes were calculated using FastSpar, n = 67 patients. All significant correlations with P ≤ 0.001 were included. Edges between nodes represent correlations. Red and blue colors indicate positive and negative correlations, respectively. Node size indicates the average abundance of the genomes. The layout of the nodes and edges was determined by Edge-weighted Spring Embedded Layout with correlation efficient as weight. (B) The distribution of different types of correlations of the genome pairs during the trial. The 3 letters show the correlations of the genome pairs at M0, M3 and M15 subsequently. Stable correlations, NNN and PPP, were highlighted (C) Average clustering of the 141 nodes based on their robust positive and negative correlations showed two clusters (green and purple range). The bar plots show the abundance changes of each node throughout the trial, which is expressed as median abundance with Z-score transformation. The differences of each node over time were tested using the Friedman test followed by Nemenyi post-hoc test. P < 0.05 was considered as significant. This panel was plotted using iTOL. (D) The seesaw-like network with the 141 nodes in two polarizing clusters. Edges between nodes represent correlations. Red and blue colors indicate positive and negative correlations, respectively. For (C) and (D), the color of the node represents the members in the two guilds: green for Guild 1 and purple for Guild 2.

We considered two genomes are connected with robust and stable ecological relationship if they keep the same type of correlation across all the three timepoints. Out of the 113,526 possible genome pairs, 92.39% showed no correlations at any of the three time points, suggesting that it may be a rare event for two genomes to establish an ecological relationship even transiently (Fig. 2B). Interestingly, 517 genome pairs showed positive correlations and 118 negative correlations at all the three time points. All the 635 stable correlations involved 184 genomes. Among these 184 HQMAGs, 43 were excluded from subsequent analysis because they had no interactions with the remaining 141 nodes (Fig. S6) and most of them showed no response to the high fiber intervention (M0 vs M3: P > 0.05, Friedman test followed by Nemenyi post-hoc). The remaining 141 HQMAGs, which included genome pairs with 468 positive and 118 negative stable correlations throughout the trial were further defined as genomes with stable ecological interactions (GSEIs) and became our microbiome signature candidates. We then explored how these 141 GSEIs were connected with each other and with the rest of the nodes in ***G***_M0_, ***G***_M3_, and ***G***_M15_ (Fig. S7A). The 141 GSEIs had significantly higher degree, betweenness centrality, eigenvector centrality, closeness centrality and stress centrality than the rest of the genomes in the networks (Fig. S7B-F). This finding indicates that these GSEIs exerted a relatively large amount of control over the interaction of other nodes (reflected by betweenness centrality and eigenvector centrality) and the information flow in the network (reflected by closeness centrality and stress centrality). Removing these GSEIs would lead to the collapse of the networks since on average 86.08% of the total edges would have been lost. These suggest that the 141 GSEIs can be considered as the core nodes of the networks as they were highly connected not only within themselves but also with other nodes.

These 141 GSEIs were also highly prevalent among participants, as 140 of them were in > 90%, and 104 were in 100% of the 74 individuals in the W group (Fig. S8). In addition, most of these 141 GSEIs were also predominant members of the gut microbiota as the abundance of 111 of them was higher than the median of the 1,845 HQMAGs and accounted for 20.78% of the total sequencing reads. Based on Bray-Curtis distance, beta-diversity analysis showed significant correlations between the profiles of the 141 GSEIs and all the 1,845 HQMAGs, as evidenced in the Mantel test (R^2^ = 0.62, P = 0.001) and Procrustes analysis (P = 0.001) (Fig. S9, Fig. 1C, D). These indicate that the variations of the 141 GSEIs contributed to the major variations of the whole gut microbial community across the 3 time points.

Based on the average linkage of these robust correlations, the 141 GSEIs fell into two clusters. There were 50 genomes in cluster 1 and 91 genomes in cluster 2 (Fig.2C). These two clusters can be considered as guilds as genome in each cluster were highly interconnected with both robust and transient positive correlations^3^ (Fig. 2D and Fig. S10). The two guilds were connected by negative edges only, indicating a competitive relationship that structures a seesaw-like network. Our data showed that the two competing guilds of the 141 GESIs existed in all three ecological networks ***G***_M0_, ***G***_M3_, and ***G***_M15_ in the W group. Furthermore, the finding of the two competing guilds in the W group at M0 suggests that such microbial organization exists irrespective of the high fiber intervention in our study. Given the similar overall gut microbiota structure between the W and U groups at M0 and in the U group across 3 timepoints (Fig. 1C, D), we speculated that two competing guilds can also be observed in the U group across the trial. Thus, we constructed the co-abundance networks based on the abundance of the 141 GESIs across the individuals in the U group at each time point. In the co-abundance networks, 99.8%, 99.51% and 99.74% of the total edges agreed with our two competing guilds, i.e., positive correlations in the guilds and negative between guilds (Fig. S11A). This showed that the detection of these two competing guilds was independent of the high fiber intervention, indicating that the such a pattern may be an inherent structure of the gut microbiome in our study.

### Functionality of the metagenomes of the two competing guilds associates with host metabolic phenotypes

We sought to determine whether the balance between the two competing guilds could be modulated by dietary fiber and describe how the two competing guilds affects the host metabolic phenotypes. In the W group, the total abundance of Guild 1 increased and Guild 2 decreased significantly from M0 to M3. Then at M15, Guild 1 decreased to a level similar to that at M0, and Guild 2 bounced back but remained lower than that at M0. Subsequently, from M0 to M3, high fiber intervention significantly increased the Guild 1-to-Guild 2 ratio. At one-year follow-up, the ratio significantly decreased and was not different from M0 (Fig. 3A). Neither the abundances of the 2 guilds nor their ratio was changed in the U group across the trial (Fig. S11B). These results showed that the changes of the balance between the two guilds composed of GSEIs were concomitant with changes in dietary fiber intake, overall gut microbiota and host phenotypes. To further validate our hypothesis that GSEIs may be essential to host health, we used the GSEIs as the selected features and applied machine leaning algorithms to explore the associations between GSEIs and each host bio-clinical parameter. Random Forest regression via leave-one-out cross-validation based on the 141 GSEIs showed that 41 out of the 43 bio-clinical parameters had significant Pearson’s correlation coefficient ranged from 0.11 to 0.44 (adjusted P value < 0.05) between the predicted and measured values (Fig. 3B). These results showed that the 141 genomes, as two competing guilds in a seesaw-like network, constitute an important microbiome signature for T2DM and the related metabolic phenotypes.

**Fig. 3.**
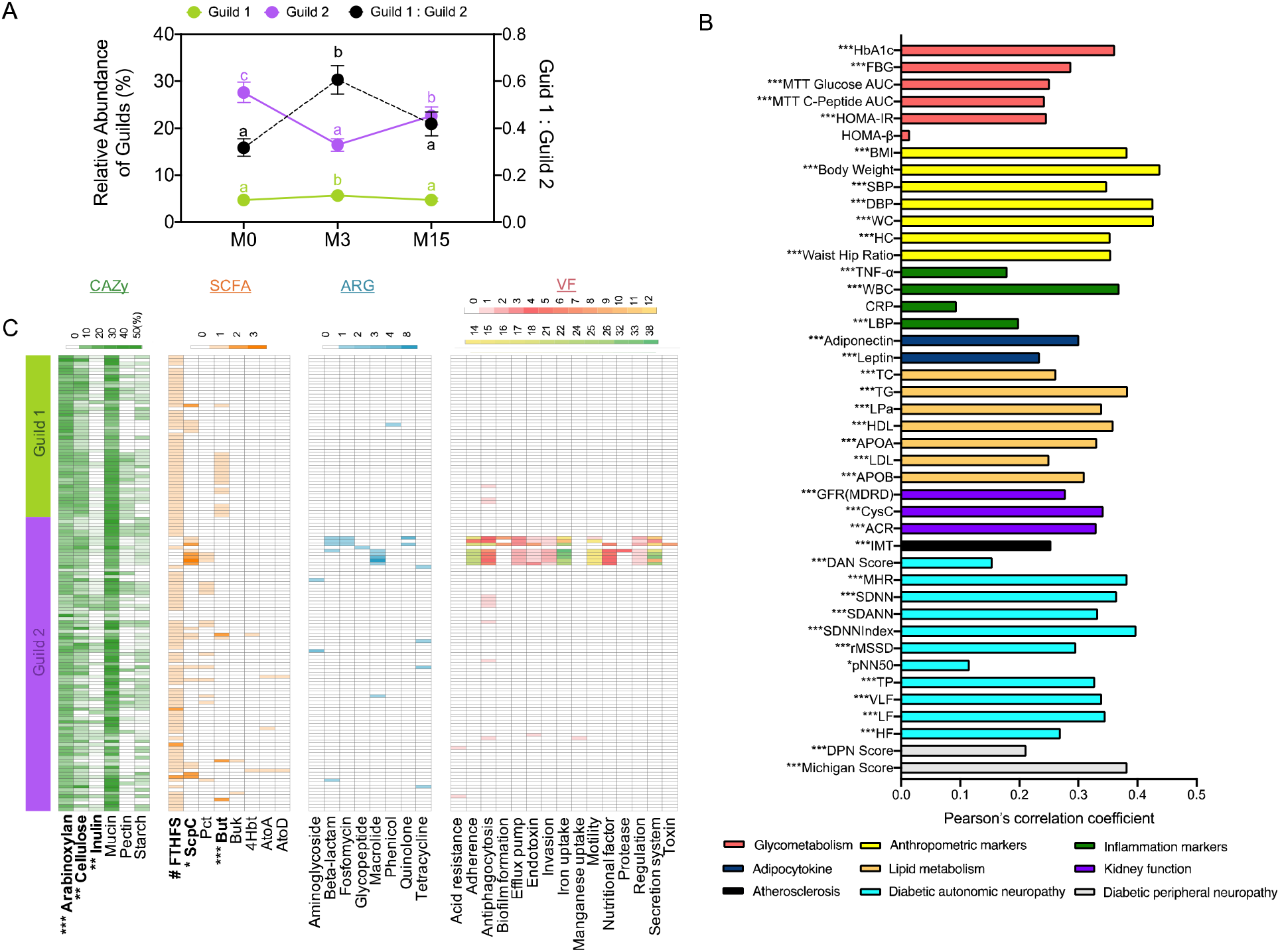
The balance between the two competing guilds in the seesaw-like network was associated with the metabolic health of patients with type 2 diabetes. (A) Change of the total abundance of Guild 1, Guild 2, and their ratio across the trial in the W group. Friedman test followed by Nemenyi test was used to analyze the difference between time points. Compact letters reflect the significance at P < 0.05. (B) Random Forest regression with leave-one-out cross-validation was used to explore the associations between the 141 genomes and the clinical parameters. The bar plot shows the Pearson’s correlations coefficient between the predicted and measured values. The asterisk before the parameter’s name shows the significance of the Pearson’s correlations. P values were adjusted by Benjamini & Hochberg’s method. * adjusted P < 0.05, ** adjusted P < 0.01 and *** adjusted P < 0.001. BMI, body mass index; SBP, systolic blood pressure; DBP, diastolic blood pressure; WC, waist circumference; HP, hip circumference; TNF-α, tumor necrosis factor-α; WBC, white blood cell count; CRP, C-reactive protein; LBP, lipopolysaccharide-binding protein; TC, total cholesterol; TG, triglyceride; Lpa, lipoprotein a; HDL, high-density lipoprotein; APOA, apolipoprotein A; LDL, low-density lipoprotein; APOB, apolipoprotein B; GFR (MDRR), glomerular filtration rate; CysC, Cystatin C; ACR, urinary microalbumin to creatinine ratio; IMT, intima-media thickness; DAN, diabetic autonomic neuropathy score; MHR, mean heart rate; SDNN, standard deviation of NN intervals; SDANN, standard deviation of the average NN intervals calculated over 5 minutes; SDNNIndex, mean of standard deviation of NN intervals for 5-minute segments; rMSSD, root-mean-square of the differences of successive NN intervals; pNN50, percentage of the interval differences of successive NN intervals greater than 50 ms; TP, total power; VLF, very low frequency power; LF, low frequency power; HF, high frequency power; DPN, diabetic peripheral neuropathy score. (C) Differences in genetic capacity of carbohydrate substrate utilization (CAZy), short-chain fatty acid production (SCFA), number of antibiotic resistance genes (ARG) and number of virulence factor genes (VF). (C) The heatmaps show the proportion (CAZy) or gene copy numbers (SCFA, ARG and VF) of each category in each genome. For carbohydrate substrate utilization, CAZy genes were predicted in each genome. The proportion of CAZy genes for a particular substrate was calculated as the number of the CAZy genes involved in its utilization divided by the total number of the CAZy genes. Arabinoxylan-related CAZy families: CE1, CE2, CE4, CE6, CE7, GH10, GH11, GH115, GH43, GH51, GH67, GH3 and GH5; cellulose-related: GH1, GH44, GH48, GH8, GH9, GH3 and GH5; inulin-related: GH32 and GH91; mucin-related families: GH1, GH2, GH3, GH4, GH18, GH19, GH20, GH29, GH33, GH38, GH58, GH79, GH84, GH85, GH88, GH89, GH92, GH95, GH98, GH99, GH101, GH105, GH109, GH110, GH113, PL6, PL8, PL12, PL13 and PL21;pectin-related: CE12, CE8, GH28, PL1 and PL9; starch-related: GH13, GH31 and GH97. For short chain fatty acid production, FTHFS: formate-tetrahydrofolate ligase for acetate production; ScpC: propionyl-CoA succinate-CoA transferase and Pct: propionate-CoA transferase for propionate production; But: Butyryl–coenzyme A (butyryl-CoA): acetate CoA transferase, Buk: butyrate kinase, 4Hbt: butyryl-CoA: 4-hydroxybutyrate CoA transferase, Ato: butyryl-CoA:acetoacetate CoA transferase (AtoA: alpha subunit, AtoD: beta subunit) for butyrate production. Mann-Whitney test (two-sided) was used to analyze the difference between Guild 1 and Guild 2. # P < 0.1, * P < 0.05, ** P < 0.01 and *** P < 0.001. Guild 1 (green bar): n = 50, Guild 2 (purple bar): n = 91.

Next, we performed genome-centric analysis of the HQMAGs in the two competing guilds to explore the genetic basis underlying the association between the dynamic changes of the seesaw networked microbiome signature and relationships with the host’s metabolic phenotypes. As the balance between the two guilds can be shifted by dietary fibers, we first sought to identify carbohydrate-active enzyme (CAZy)-encoding genes and genes encoding key enzymes in short-chain fatty acid (SCFA) production to compare the genetic capacity for carbohydrate utilization between the two guilds. Compared with genomes in Guild 2, those in Guild 1 enriched CAZy genes for arabinoxylan (P < 0.001), cellulose (P < 0.01) and had lower proportion of CAZy genes for inulin utilization (P < 0.01) (Fig. 3C, Table S5). There was no difference in genes for starch, pectin, and mucin utilization between the two guilds. Our previous study showed that gut microbiota benefited patients with T2DM via acetic and butyric acid production from carbohydrate fermentation^11^. Among the terminal genes for the butyrate biosynthetic pathways from both carbohydrates (i.e., *but* and *buk*) and proteins (i.e., *atoA*/*D* and *4Hbt*), the copy number of *but* was significantly higher in Guild 1 and there was no difference in the other terminal genes between the two guilds (Fig. 3C). More than one-third of the genomes in Guild 1 harbored the *but* gene while less then 5% of the genomes in Guild 2 had this gene (Fisher’s exact test P < 0.001). Compared with Guild 2, Guild 1 also trended higher in its genetic capacity for acetate production (P = 0.06) but had a lower genetic capacity for propionate production (P < 0.05) (Fig. 3C). These results showed that compared to Guild 2, Guild 1 had significantly higher genetic capacity for utilizing complex plant polysaccharides and producing acetate and butyrate.

From the perspective of pathogenicity, 21 out of the 1,845 HQMAGs encoded 750 virulence factor (VF) genes. Among the 21 VF-encoding genomes, 3 were in Guild 1 while 18 were in Guild 2. Three out of the 50 genomes in Guild 1 had one VF gene involved in antiphagocytosis. In Guild 2, 18 out of the 91 genomes encoded 747 VF genes across 15 different VF classes i.e., acid resistance, adherence, antiphagocytosis, biofilm formation, efflux pump, endotoxin, invasion, iron uptake, manganese uptake, motility, nutritional factor, protease, regulation, secretion system, and toxin (Fig. 3C, S12A). Notably, 98.53% of all the VF genes in Guild 2 were harbored in 8 genomes (1 in *Enterobacter kobei*, 2 in *Escherichia flexneri*, 3 in *Escherichia coli* and 2 in *Klebsiella*). The highly enriched genes for virulence factors in genomes of Guild 2 (Fisher’s Exact test, P < 2.2×10^−16^,) indicates that this guild may play an important role in aggravating the metabolic disease phenotypes. In terms of antibiotic resistance genes (ARG), in Guild 1, only 1 genome (2.00% of the genomes in this guild) harbored a copy of an ARG related to phenicol (Fig. 3C, S12B). In Guild 2, 17 genomes (18.68% of the genomes in this guild) encode 40 ARGs for resistance to 7 different antibiotic classes i.e., aminoglycosides, beta-lactam, fosfomycin, glycopeptide, quinolone, macrolide, and tetracycline. Thus, Guild 2 may serve as a reservoir of ARGs for horizontal transfer to opportunistic pathogens. Taken together, our data showed that the two competing guilds had distinct genetic capacity with Guild 1 being potentially beneficial and Guild 2 detrimental.

### The seesaw networked microbiome signature exists in cohorts across ethnicity and geography

We then asked that whether these 141 genomes, organized as two competing guilds in a stable seesaw-like network, is this a microbiome signature unique in T2DM or it is common in other diseases. To answer this question, we used the 141 GSEIs in our seesaw-like network as reference genomes to perform read recruitment analysis, which is a commonly used method to estimate abundance of reference genomes^22,23^ in metagenomes (Fig. S13). In an independent T2DM study^24^, 32.92% of the reads were recruited and 128 of the GSEIs were detected as part of a co-abundance network based on their estimated abundance across the T2DM patients. In this co-abundance network, 97.82% of the total edges followed the pattern in our seesaw-like network (i.e., positive edges within each guild and negative edges between the 2 guilds) (Fig. 4A), which further supported the existence of this seesaw-like network in T2DM patients.

**Fig.4.**
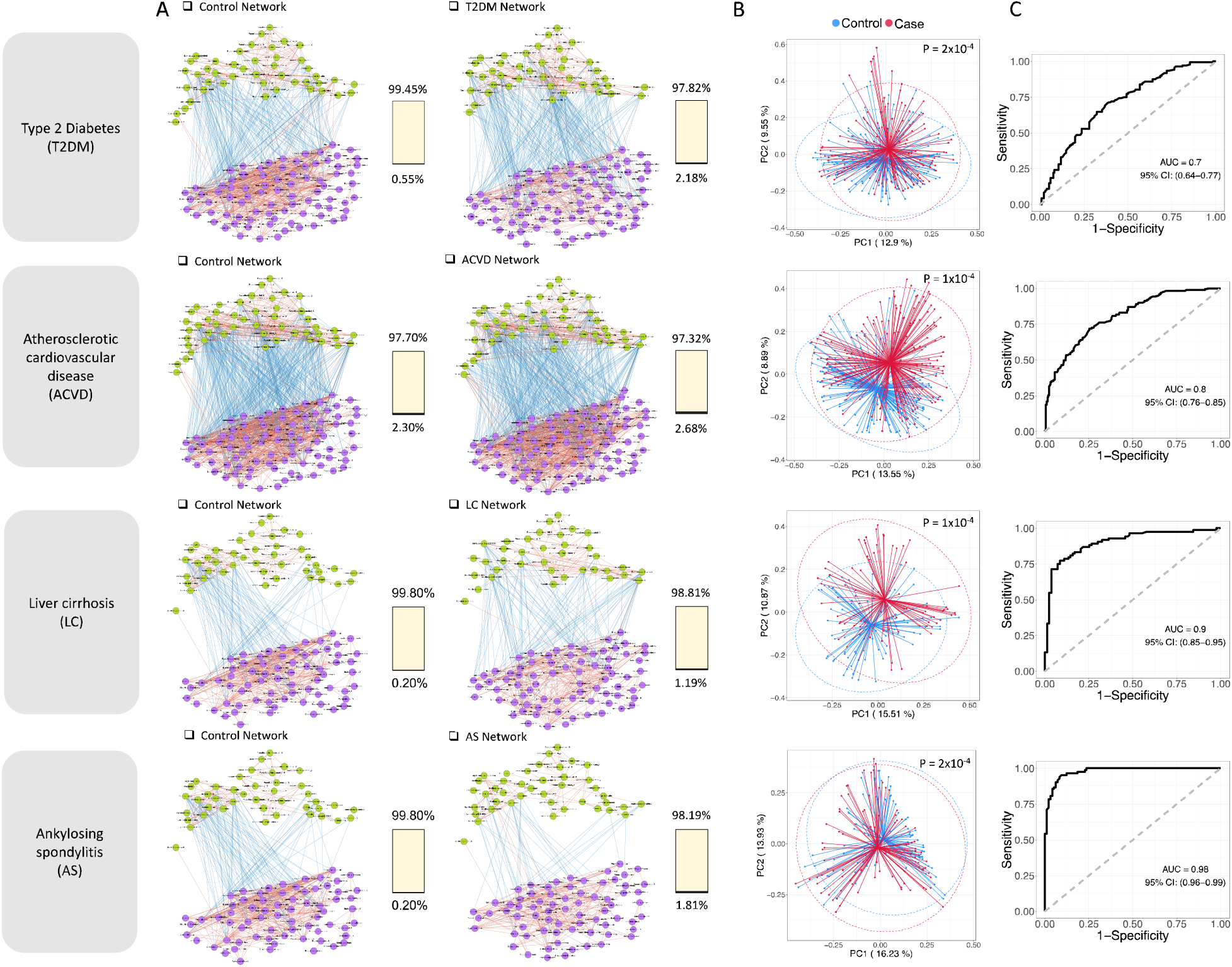
The seesaw networked microbiome signature exists in other independent human cohorts and supports classification models for different diseases. (A) Members of the two competing Guilds in the seesaw networked microbiome signature showed similar ecological interactions in four independent human gut metagenomic datasets. The correlations between the genomes were calculated using FastSpar. All significant correlations (P ≤ 0.001) were included. Lines between nodes represent correlations, and red and blue colors indicate positive and negative correlations, respectively. The color of the node represents the members in the two competing guilds: green for Guild 1 and purple for Guild 2. The percentage of correlations followed the pattern in the seesaw networked microbiome signature (i.e., positive edges within each guild, negative edges between the 2 guilds) was in yellow, and the ratio of correlations that were negative within each guild and positive between the guilds was in black of the 100% stacked bar. (B) The composition of the microbiome signature was different between control and patients in each dataset as shown in the Principal Coordinates Analysis plot based on Bray-Curtis distance. 95% confidence ellipses were projected for control and patients respectively. The p values of the PERMANOVA test were indicated. (C) The microbiome signature supports classification models for the four different diseases. The area under the ROC curve (AUC) of the Random Forest classifier based on the 141 genomes in the microbiome signature to classify control and patients in each dataset. Leave-one-out cross validation was applied. Type 2 diabetes (T2DM): Control n = 136, T2D n=136; Atherosclerotic cardiovascular disease (ACVD): Control n = 171 and ACVD n = 214; Liver cirrhosis (LC): control n = 83 and LC n =84; Ankylosing spondylitis: Control n = 83 and AS n = 97.

Moreover, 35.28% of the reads were recruited in the metagenomes of 136 healthy controls of the same study^24^ and 119 of the GSEIs were constructed into a co-abundance network in which 99.45 % of the total edges agreed with our seesaw-like network (Fig. 4A). In the context of beta-diversity based on Bray-Curtis distance, our microbiome signature showed significant differences (PERMAONVA test P = 2×10^−4^) between T2DM patients and the healthy controls based on the abundance matrix of the reference genomes (Fig. 4B). This suggests that the variation of this microbiome signature was associated with T2DM in this independent dataset. To further validate such associations, using the abundance matrix of the genomes in the microbiome signature as input features and the phenotype data as target variable, we constructed Random Forest regression models and found that this microbiome signature was significantly correlated with BMI, fasting insulin, and HbA1c (Fig S14). Furthermore, we developed a machine learning classifier based on a Random Forest algorithm to see if we can classify patients and control. Receiver operating characteristic curve analysis showed a moderate diagnostic power with area under the curve (AUC) of 0.70 by a leave-one-out cross-validation. Thus, we showed that our seesaw networked microbiome signature not only existed in an independent T2DM study but also maintained a similar relationship with the host metabolic phenotypes.

We then extended our hypothesis that the seesaw networked microbiome signature represents an inherent feature of human gut microbiome and the disruption of which may be related to diseases in addition to T2DM. We first performed the same validation analysis in metagenomic datasets of three different types of diseases, including atherosclerotic cardiovascular disease (ACVD) ^25^, liver cirrhosis (LC)^26^ and ankylosing spondylitis (AS)^27^. In ACVD patients and their controls, 36.21% and 32.73% of the reads were recruited respectively, and 134 genomes from the patients and 133 genomes from the controls were constructed into co-abundance networks with 97.32% and 97.70% of the total edges respectively agreed with our seesaw-like network (Fig. 4A). There were 33.84%, 35.83% and 41.02% of the reads recruited to the reference genomes in the metagenomic datasets of the healthy control (the studies on LC and AS employed the same control cohort), LC and AS patients respectively. With the 141 GSEIs as reference, 112, 113 and 113 genomes were constructed into co-abundance networks with 99.80%, 98.81% and 98.19% of the total edges agreed with our seesaw-like network in the metagenomic datasets of the healthy control, LC and AS patients respectively (Fig. 4A). In the PCoA plot based on Bray-Curtis distance, our microbiome signature showed significant differences (PERMANOVA test, P < 0.001) between control and patients in all 3 datasets (Fig. 4B). In the LC study, we also used the abundance matrix of the genomes in the microbiome signature as input features and the phenotype data as target variable to construct Random Forest regression models and found that our microbiome signature was significantly correlated with total bilirubin, albumin level, and BMI (Fig. S15). Compared with the T2DM dataset^24^, the Random Forest classifier based on our microbiome signature showed better diagnostic power in distinguishing case from control for ACVD (AUC = 0.80), LC (AUC = 0.90), and AS (AUC = 0.98) (Fig. 4C).

To further confirm the relevance of this microbiome signature to human diseases, we estimated the abundances of the genomes from this microbiome signature in datasets from more disease types and across different ethnicity and geography. These datasets included inflammatory bowel diseases (IBD) (American cohort and Dutch cohort), colorectal cancer (CRC) (Chinese cohort, Australian cohort and German cohort), schizophrenia (Chinese cohort), and Parkinson’s disease (PD) (Chinese cohort). On average, 31.32% ± 4.21% (mean ± SD) of reads were recruited to the reference genomes in these datasets. We validated that this microbiome signature showed diagnostic power to classify case and control in the metagenomic dataset from studies on IBD (AUC = 0.71 for IBD dataset 1^28^, AUC=0.91 for IBD dataset 2^29^ and AUC=0.83 for IBD dataset 3^29^), CRC (AUC = 0.74 for CRC dataset 1^30^, AUC = 0.75 for CRC dataset 2^31^ and AUC = 0.71 for CRC dataset 3^32^), schizophrenia^33^ (AUC = 0.68), and PD^34^ (AUC = 0.77) (Fig. S16). In addition, we used MMUPHin^35^ to correct batch effects from the different cohorts in IBD and CRC and applied leave-one-cohort-out (LOCO) analysis^36^ to evaluate the universality of the diagnostic power of this microbiome signature in these two diseases. The AUC values from LOCO analysis were 0.77 to 0.84 for IBD and 0.68 to 0.70 for CRC (Fig. S17). These results showed the existence of our microbiome signature in healthy controls and various patient populations across ethnicity and geography from independent studies. The associations between the 141 genomes and host phenotypes and their discriminative power as biomarkers to classify controls vs. patients with various types of diseases indicate that these genomes, organized as two guilds in a seesaw-like network, represent a common microbiome signature associated with a wide range of human diseases.

## Discussion

In the current study, our genome-centric, reference-free, and ecological-interaction-focused approach led to the identification of a robust seesaw-like network of two competing guilds of bacterial genomes, whose changes were associated with a wide range of host phenotypes in patients with T2DM. Moreover, random forest models based on these genomes classified case and control across a wide range of diseases, indicating that these genomes may form a core microbiome signature that exists in populations of diverse ethnicity, geography, and disease status.

Our novel microbiome signature organizes genomes in a seesaw-like network exhibiting both cooperative and competitive interactions. Cooperative ecological networks are expected to promote overall metabolic efficiency, such as the co-operative metabolism that benefits the host^37^. However, they also create dependency and the potential for mutual downfall that may bring destabilizing effect on the gut microbial ecosystem. This destabilizing effect of cooperation can be dampened by introducing ecological competition into the network^37^. Thus, a seesaw-like network with both cooperative and competitive interactions may represent the key characteristic of a stable microbiome structure^37^. Interestingly, while the seesaw-like network is stable, the weight of the two ends i.e., the abundances of Guild 1 and Guild 2, are modifiable and such changes are associated with host health. When large amount of complex fiber was introduced or withdrawn, Guilds 1 and 2 showed no change in membership nor in the types of interactions with each other but experienced dramatic shifts in guild-level abundance in a competing manner. Members in Guild 1 have higher genetic capacity for degrading complex plant polysaccharides and produce beneficial metabolites including SCFAs which may suppress populations of pathobionts in Guild 2^11^. Members of Guild 2 need to be kept low since their overgrowth may jeopardize host health by increasing inflammation, etc.^38^. However, pathobionts in Guild 2 cannot be eliminated, e.g., they could serve as the necessary agents that train our immune system from early days^39,40^. Therefore, the balance between Guild 1 and Guild 2 becomes critical in determining whether the gut microbiome supports health or aggravate diseases. This seesaw-like network between Guilds 1 and 2 allows the genomes in our microbiome signature to readily respond to changes of external energy input to the gut microbial ecosystem and mediate its impact on host health, while simultaneously maintains its structural integrity. Such structural integrity may be key to long-term ecological stability of the gut microbiome and its ability to provide essential health-relevant functions to the host.

The fact that this seesaw-like network can be detected in other independent metagenomic datasets and is shown correlated with different diseases indicates that such an ecological organization is not specific to T2DM nor dependent on the high fiber diet we used, Such a seesaw networked structure may have been stabilized by natural selection over a long history of co-evolution between microbiomes and their hosts^16 41^. A selection pressure may have been exerted by dietary fibers that interact directly with gut bacteria as external energy source^42,43^. Studies on coprolites showed that dietary fiber intake was much higher in ancient humans and only reduced significantly in the past 150 years^44,45^ (130 g/d of plant fiber intake in prehistoric diet^46^ vs. a median intake of 12–14 g/d in the modern American diet^47^). Such a high fiber intake over evolutionary history may have favored beneficial bacteria in Guild 1 due to their higher genetic capacity to utilize plant polysaccharides as an external energy supply, enabling them to gain competitive advantage over pathobionts in Guild 2^48^. Akin to tall trees as the foundation species for a closed forest, Guild 1 may work as the “foundation guild” for stabilizing a healthy gut microbiome and keeping the pathobionts at bay^49^. The dominance of Guild 1 over Guild 2 can increase host fitness as shown by the epidemiologically and clinically proven health benefits of dietary fibers in both preventing and alleviating a wide range of chronic conditions^11,43,50,51^.

Moreover, the two seesaw networked guilds may constitute a core gut microbiome signature in humans^52,53^, since 1) they are commonly shared among populations across ethnicity and geography; 2) they show temporal stability not only in membership but also in their interactions with each other and the host; 3) they make up about 10% of the gut microbiome membership but are disproportionally important for shaping the ecological community; 4) they support the provision of essential health-relevant functions to the host; and 5) such a core microbiome organized in a seesaw-like network may have been established over a long history of co-evolution.

As an evolutionarily conserved ecological structure, our seesaw-like network demonstrated remarkably stable relationships both internally within the network and externally with the host. Within the seesaw-like network, it is the imbalance between the two competing guilds that may play a role as the common biological basis for many human diseases. Targeting this core microbiome signature to restore and maintain ecological dominance of the beneficial guild over the detrimental guild could be fundamentally important to human health recovery and maintenance.

## Materials and Methods

### Clinical Experiment

#### Study design^11^

This clinical trial, conducted at the Qidong People’s Hospital (Jiangsu, China), examined the effect of a high fiber diet in free-living conditions in a cohort of individuals clinically diagnosed T2DM (QIDONG). The study protocol was approved by Ethics Committee of Shanghai General Hospital (2014KY104), and the study was conducted in accordance with the principles of the Declaration of Helsinki. All participants provided written informed consent. The trial was registered in the Chinese Clinical Trial Registry (ChiCTR-IPC-14005346). The study design and participant flow are shown in Fig. S1.

T2DM patients of the Chinese Han ethnicity were recruited for the study (age: 37 - 70 years; HbA1c: 6.5% - 12.0%. More detailed description of inclusion and exclusion criteria were shown in Chinese Clinical Trial registry (http://www.chictr.org.cn).

Patients received either a high-fiber diet (WTP diet) as the treatment group (W group) or the usual care (Usual diet) as the control group (U group) for 3 months. Total caloric and macronutrients prescriptions were based on age-specific Chinese Dietary Reference Intakes (Chinese Nutrition Society, 2013). The WTP diet, based on wholegrains, traditional Chinese medicinal foods and prebiotics, included three ready-to-consume pre-prepared foods^11^. The usual care included standard dietary and exercise advice that was made according to the Chinese Diabetes Society guidelines for T2DM^54^. Patients in W group were provided with the WTP diet to perform a self-administered intervention at home for three months, while patients in U group accepted the usual care. W group stopped WTP diet intervention at the end of the third month (at M3). Then W and U continued a one-year follow-up (M15). A meal-based food frequency questionnaire and 24-h dietary recall were used to calculate nutrient intake based on the China Food Composition 2009^55^. Patients in both groups continued with their antidiabetic medications according to their physician prescriptions (Table S1).

Before a 2-week run-in period, all participants attended a lecture on diabetes intervention and improvements and received diabetes education and metabolic assessments. 119 eligible individuals were enrolled based on the inclusion and exclusion criteria and assigned into two groups in a 2:1 ratio (n = 79 in W group, n = 40 in U group) determined by SAS software.

Physical examinations were carried out at M0, M3, and M15 in Qidong People’s Hospital (Jiangsu, China). Sample collection instructions were provided to the participants at the day before. The participants provided the feces and first early morning urine as requested. After collecting fasting venous blood sample, a 3-h meal tolerance test (Chinese buns containing 75 g of available carbohydrates; MTT test) was conducted and the postprandial venous blood samples at 30, 60, 120, and 180 min were collected. All the blood samples were centrifuged at 3000 rpm for 20 min at 4◻ after standing at room temperature for 30 min to obtain serum. The fasting blood serum were divided into two parts, one used for hospital tests and the other used for lab tests. The feces, urine, and serum samples were stored in dry ice immediately then transported to lab and frozen at −80°C. Subsequently, anthropometric markers and diabetic complication indexes were measured. Ewing test^56^ and 24-h dynamic electrocardiogram were conducted to estimate diabetic autonomic neuropathy (DAN). B-mode carotid ultrasound was conducted to estimate atherosclerosis. Michigan Neuropathy Screening Instrument^57^ was conducted to estimate diabetic peripheral neuropathy (DPN). In addition, A meal-based food frequency questionnaire and the 24-h dietary review were recorded for nutrient intake calculation. The drug use was self-reported and presented in table S1.

The fasting venous blood was used to measure HbA1c, fasting blood glucose, fasting insulin, fasting C-Peptide, C-reactive protein (CRP), blood routine examination, blood biochemical examination and five analytes of thyroid. The venous blood samples at 30, 60, 120, and 180 min of MTT were used to measure the postprandial blood glucose, insulin, and C-Peptide. The fasting early morning urine was used to measure the routine urine examination and urinary microalbumin creatinine ratio. The measurements above were completed at Qidong People’s Hospital. Fasting venous blood was used to quantify TNF-α (R&D Systems, MN, USA), lipopolysaccharide-binding protein (Hycult Biotech, PA, USA), leptin (P&C, PCDBH0287, China) and adiponectin (P&C, PCDBH0016, China) by enzyme-linked immunosorbent assays (ELISAs) at Shanghai Jiao Tong University.

The homeostatic model assessments of insulin resistance (HOMA-IR) and islet β-cell function (HOMA-β) were calculated based on fasting blood glucose (mmol/L) and fasting C-Peptide (pmol/L)^58^: HOMA-IR = 1.5 + FBG * Fasting-C-Peptide / 2800; HOMA-β = 0.27 * Fasting-C-Peptide / (FBG - 3.5). Glomerular Filtration Rate was estimated by formula GFR (ml/min per 1.73 m^2^) = 186 * Scr^−1.154^ * age^−0.203^ * 0.742 (if female) * 1.233 (if Chinese)^59^, where Scr (serum creatinine) is in mg/dl and age is in years.

### Gut microbiome analysis

#### Metagenomic sequencing

DNA was extracted from fecal samples using the methods as previously described^10^. Metagenomic sequencing was performed using Illumina Hiseq 3000 at GENEWIZ Co. (Beijing, China). Cluster generation, template hybridization, isothermal amplification, linearization, and blocking denaturing and hybridization of the sequencing primers were performed according to the workflow specified by the service provider. Libraries were constructed with an insert size of approximately 500 bp followed by high-throughput sequencing to obtain paired-end reads with 150 bp in the forward and reverse directions. Table S3 shows the number of raw reads of each sample.

#### Data quality control

Prinseq^60^ was used to: 1) trim the reads from the 3′ end until reaching the first nucleotide with a quality threshold of 20; 2) remove read pairs when either read was < 60 bp or contained “N” bases; and 3) de-duplicate the reads. Reads that could be aligned to the human genome (H. sapiens, UCSC hg19) were removed (aligned with Bowtie2^61^ using --reorder --no-hd --no-contain --dovetail). Table S3 shows the number of high-quality reads of each sample for further analysis.

#### De novo assembly, abundance calculation, and taxonomic assignment of genomes

De novo assembly was performed for each sample by using IDBA_UD^62^ (--step 20 --mink 20 --maxk 100 --min_contig 500 --pre_correction). The assembled contigs were further binned using MetaBAT^63^ (--minContig 1500 --superspecific -B 20). The quality of the bins was assessed using CheckM^64^. Bins had completeness > 95%, contamination < 5% and strain heterogeneity < 5% were retained as high-quality draft genomes (Table S6). The assembled high-quality draft genomes were further dereplicated by using dRep^65^. DiTASiC^66^, which applied kallisto for pseudo-alignment^67^ and a generalized linear model for resolving shared reads among genomes, was used to calculate the abundance of the genomes in each sample, estimated counts with P-value > 0.05 were removed, and all samples were downsized to 36 million reads (One sample with read mapping ratio < 25%, which could not be well represented by the high quality genomes, were removed in downstream analysis). Taxonomic assignment of the genomes was performed by using GTDB-Tk^68^ with default parameters (Table S7).

#### Gut microbiome functional analysis

Prokka^69^ was used to annotate the genomes. KEGG Orthologue (KO) IDs were assigned to the predicted protein sequences in each genome by HMMSEARCH against KOfam using KofamKOALA with default parameters^70^. Antibiotic resistance genes were predicted using ResFinder^71^ with default parameters. The identification of virulence factors were based on the core set of Virulence Factors of Pathogenic Bacteria Database (VFDB^72^, download July 2020). The predicted proteins sequences were aligned to the reference sequence in VFDB using BLASTP (best hist with E-value < 1e-5, identity > 80% and query coverage > 70%). Genes encoding carbohydrate-active enzymes (CAZys) were identified using dbCAN (releasee 6.0)^73^, and the best-hit alignment was retained. Genes encoding formate-tetrahydrofolate ligase, propionyl-CoA: succinate-CoA transferase, propionate CoA-transferase, 4Hbt, AtoA, AtoD, Buk and But were identified as described previously^11^.

#### Gut microbiome network construction and analysis

In W group, prevalent genomes shared by more than 75% of the samples at every timepoint were used to construct the co-abundance network at each timepoint. Fastspar^74^, a rapid and scalable correlation estimation tool for microbiome study, was used to calculate the correlations between the genomes with 1,000 permutations at each time point based on the abundances of the genomes across the patients and the correlations with P ≤ 0.001 were retained for further analysis. The networks were visualized with Cytoscape v3.8.1^75^. The layout of the nodes and edges was determined by Edge-weighted Spring Embedded Layout using the correlation coefficient as weights. The links between the nodes are treated as metal springs attached to the pair of nodes. The correlation coefficient was used to determine the repulsion and attraction of the spring^75^. The layout algorithm sets the position of the nodes to minimize the sum of forces in the network. We defined robust stable edges as the unchanged positive/negative correlations between the same two genomes across all the 3 networks at M0, M3, and M15. Stable genome pairs were clustered based on robust positive (set as 1) and negative (set as −1) edges with average clustering. We used iTOL^76^, an online tool for display, manipulation, and annotation for various trees, to integrate and visualize the clustering tree, taxonomy information, and abundance changes of the 141 genomes.

#### Validation in independent cohorts

Twelve independent metagenomic datasets were downloaded from SRA or ENA database. The group information was collected from the corresponding papers or from curatedMetagenomicData^77^ (Table S8). DiTASiC was used to recruit reads and estimate the abundance of the 141 genomes in each sample, estimated counts with P-value > 0.05 were removed and further converted to relative abundance divided by the total number of reads. To reduce false positive in the validation dataset, relative abundance < 0.001% were further removed. A random forest classification model to classify case and control was constructed based on the estimated abundances of the genomes in each dataset with leave-one-out cross-validation.

MMUPHin^35^ was used to adjust the estimated abundances of the genomes by correcting batched effects from the different cohorts in IBD and CRC studies. Random forest classification models with leave-one-cohort-out analysis were further performed on the adjusted abundance matrix^36^.

Datasets from 4 studies were included to validate the commonality of the seesaw-like network. These datasets were from 136 control and 136 T2DM individuals in Qin et al., 2012^24^; 171 control and 214 atherosclerotic cardiovascular disease individuals in Jie et al., 2017^25^; 83 control and 84 liver cirrhosis individuals in Qin et al., 2014^26^; and 83 control and 97 ankylosing spondylitis individuals in Wen et al., 2017^27^. Fastspar was used to calculate the correlations between the genomes with 1,000 permutations and the correlations with P ≤ 0.001 were remained for constructing the networks. 30 repeat 5-fold cross-validation was used and the correlations shared by more than 95% of the 150 networks constructed from the cross-validation process were remained in the final network.

### Statistical Analysis

Statistical analysis was performed in the R environment (R version3.6.1). Friedman test followed by Nemenyi post-hoc test was used for intra-group comparisons for repeat measurements. Mann-Whitney test (two-sided) was used for comparisons between W and U at the same time point. Pearson Chi-square tests was performed to compare the differences of categorical data between groups or timepoints. PERMANOVA test (9,999 permutations) was used to compare the groups of gut microbiota structure.

Mann-Whitney test (two-sided) and Fisher’s exact test (two sided) were used to compare the functions between Guild 1 and Guild 2. Random Forest with leave-one-out cross-validation was used to perform regression and classification analysis based on this microbiome signature and clinical parameters/groups.

## Acknowledgements

This work was supported by grants from National Natural Science Foundation of China (31930022, 81871091, 81870594 and 81870596), the National Key Research and Development Project (2019YFA0905600), the School of Environmental and Biological Sciences and the New Jersey Institute for Food, Nutrition, and Health (seed pilot grant cycle 1, awarded in 2019), Canadian Institute for Advanced Research, Notitia Biotechnologies Company, Clinical Research Plan of SHDC [No. SHDC2020CR1016B], and the Project of Songjiang District Municipal Health Commission (0702N18003).

## Author contributions

L.Z., Y.P., C.Z. and Y.S. designed the supervised the study; T.X., F.Z., X.D. and Y.S. managed the clinical research; T.X., X.D., D.W., J.F., Y.S., X.L., S.J., X.W. and F.Z. conducted the clinical study; T.X., X.W., H.F. and C.Z. performed clinical testing and data analysis; H.F. provided dietary counselling to the participants and conducted dietary analysis; T.X. and X.W. prepared the fecal DNA for sequencing. G.W., X.T., C.Z. performed microbial data analysis. G.W. performed data validation in independent datasets. L.Z., C.Z., G.W., N.Z. and Y.Y.L. wrote and revised manuscript. G.W., T.X., N.Z., Y.Y.L. and X.D. contributed equally to this work.

## Data availability

The metagenomic sequencing data has been deposited under accession numbers PRJEB15179 and PRJEB52732

## Code availability

Parameters of the bioinformatic tools applied in the study were showed in the method section. Scripts and command lines related to the current study are freely available from the corresponding author (L.Z.) upon request.

## Ethics & Inclusion statement

We have carefully considered research contributions and authorship criteria when involved in multi-region collaborations involving local researchers so as to promote greater equity in research collaborations.

