## Supplemental Figures and Tables for "Two Competing Guilds as a Core Microbiome Signature for Health Recovery"

### Supplementary figures

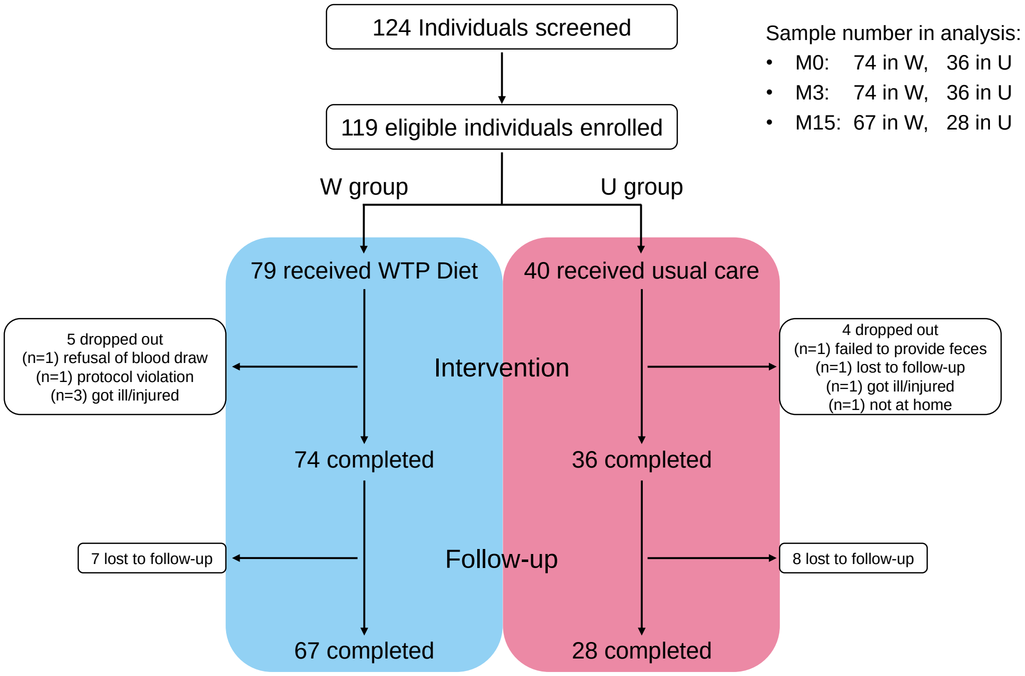


**Fig. S1. The flow diagram of participants in the trial.**

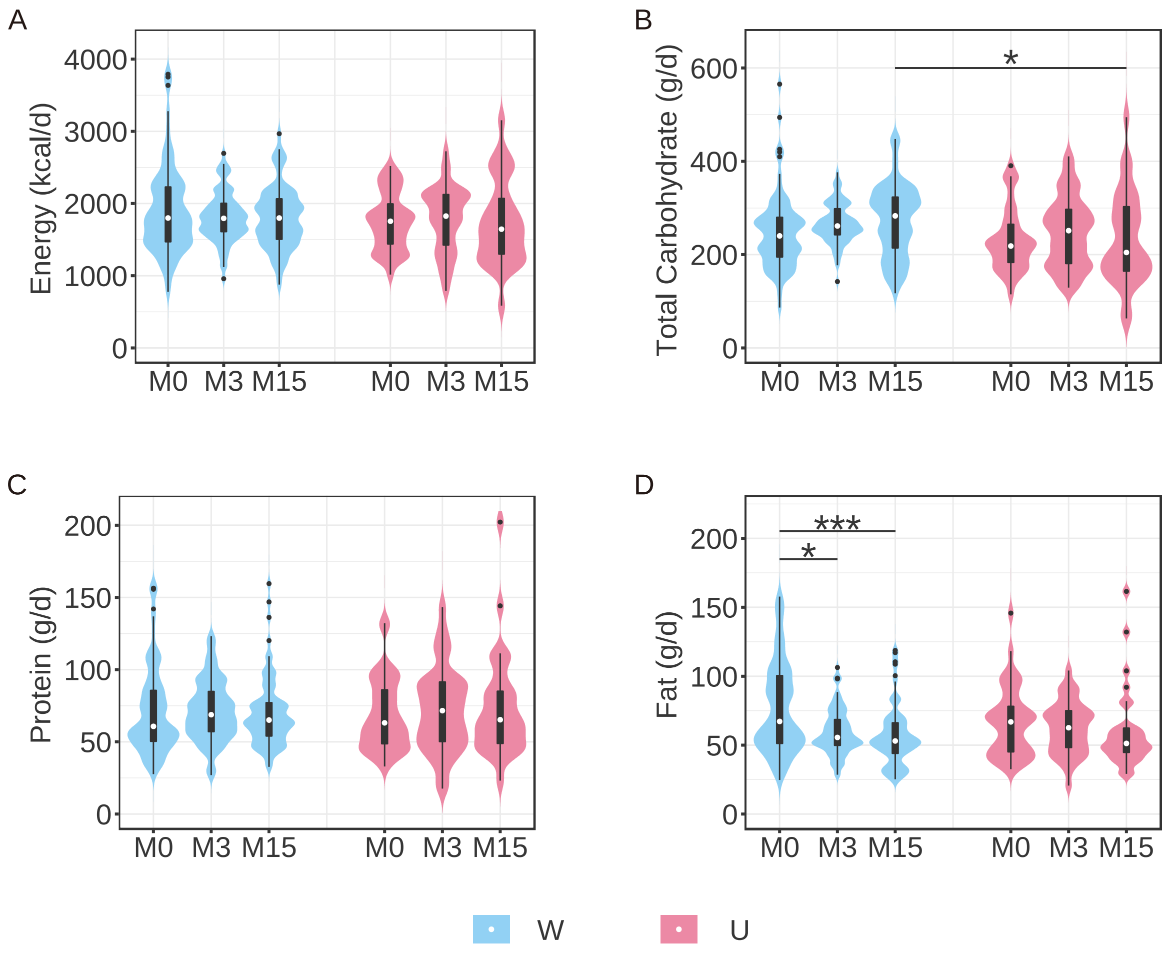


**Fig.S2. Violin plot of energy and macronutrient intake during the trial in W and U group**. Friedman test followed by Nemenyi post-hoc test was used for comparison in the same group. Mann-Whitney test (two-sided) was used for comparison between W and U at the same time point. *P < 0.05, **P < 0.01 and ***P < 0.001. Boxes show the medians and the interquartile ranges (IQRs), the whiskers denote the lowest and highest values that were within 1.5 times the IQR from the first and third quartiles, and outliers are shown as individual points.

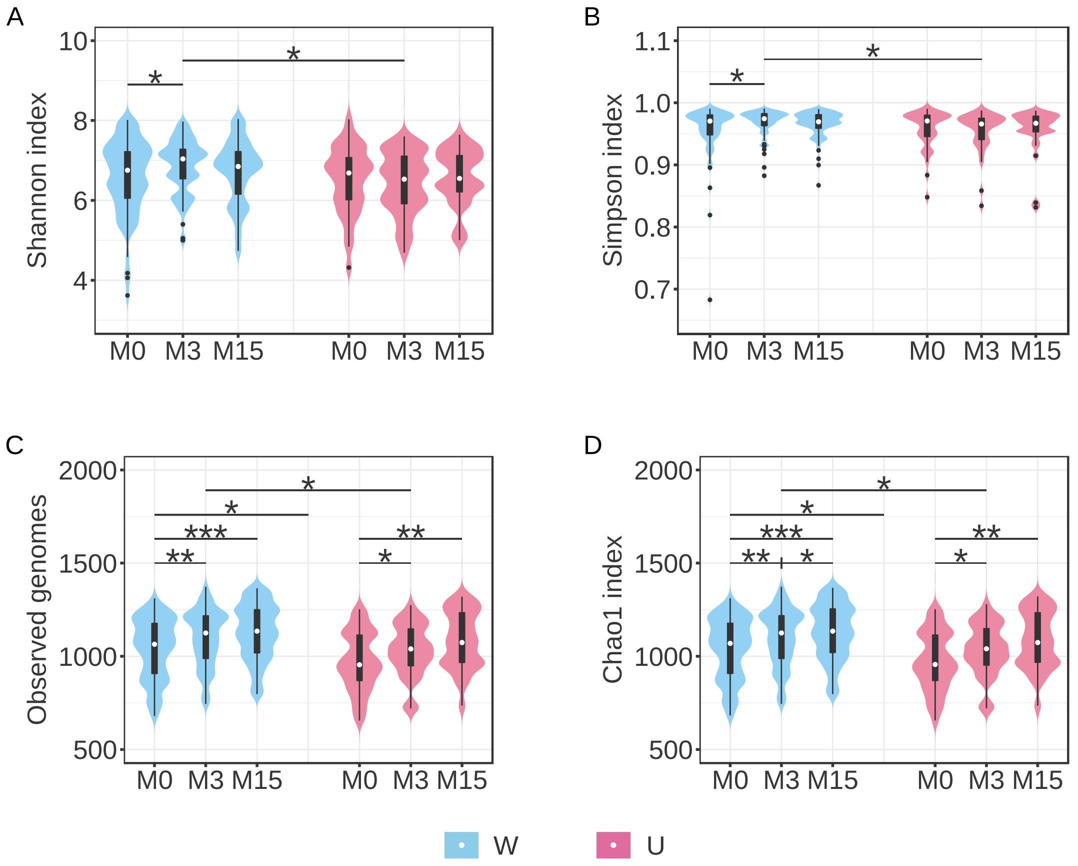


**Fig. S3 Violin plot of the change in alpha diversity of gut microbiomes during the trial in W and U group.** (A) Shannon Index; (B) Simpson Index; (C) Observed Genomes; (D) Chao 1 Index. Friedman test followed by Nemenyi post-hoc test was used for comparison in the same group. Mann-Whitney test (two-sided) was used for comparison between W and U at the same time point. *P < 0.05, **P < 0.01 and ***P < 0.001. Boxes show the medians and the interquartile ranges (IQRs), the whiskers denote the lowest and highest values that were within 1.5 times the IQR from the first and third quartiles, and outliers are shown as individual points.

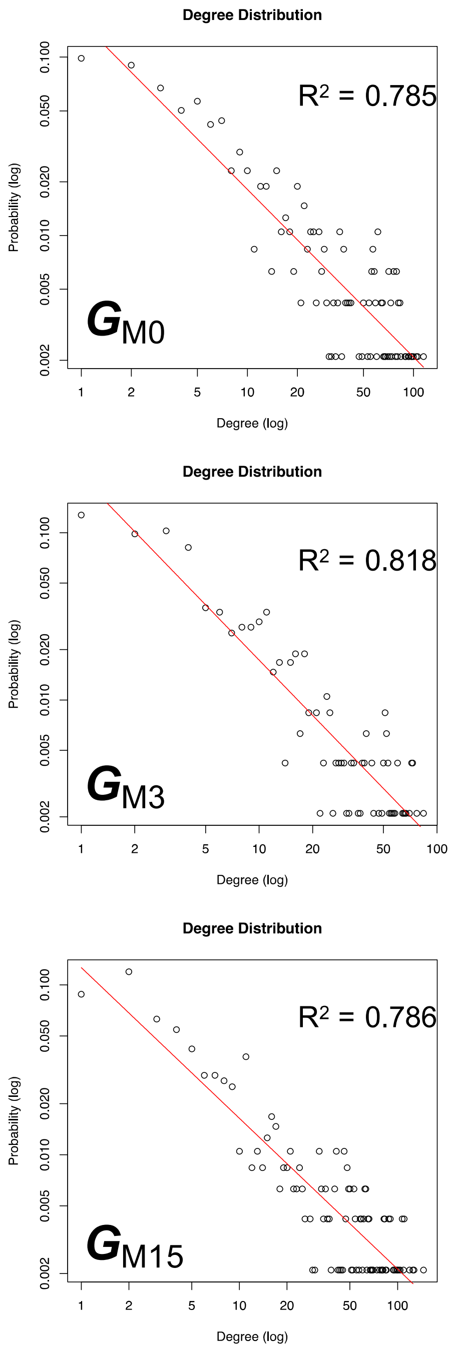


**Fig S4. The co-abundance networks of the prevalent genomes were scale-free networks across the trial**. Degree distribution were fitted well with power law model.

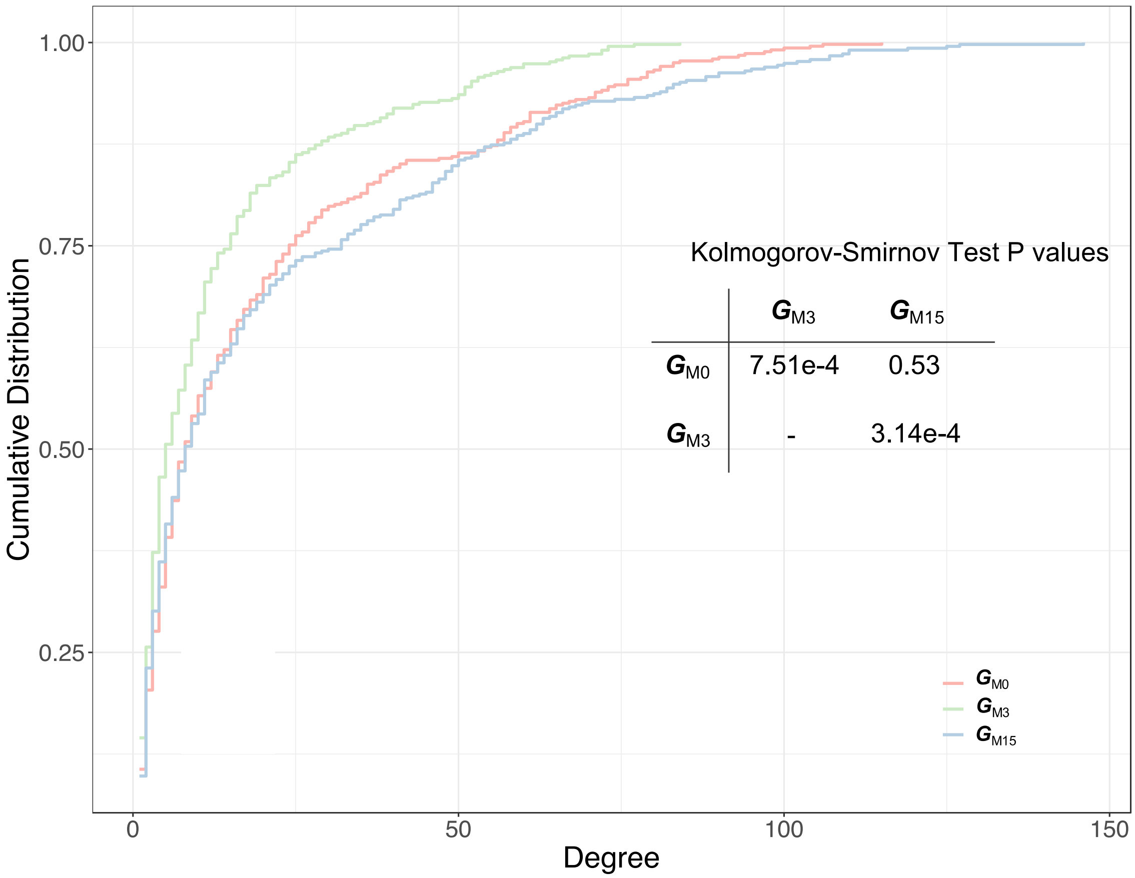


**Fig. S5. Introduction and withdrawal of high fiber intervention significantly change the network degree distribution.**

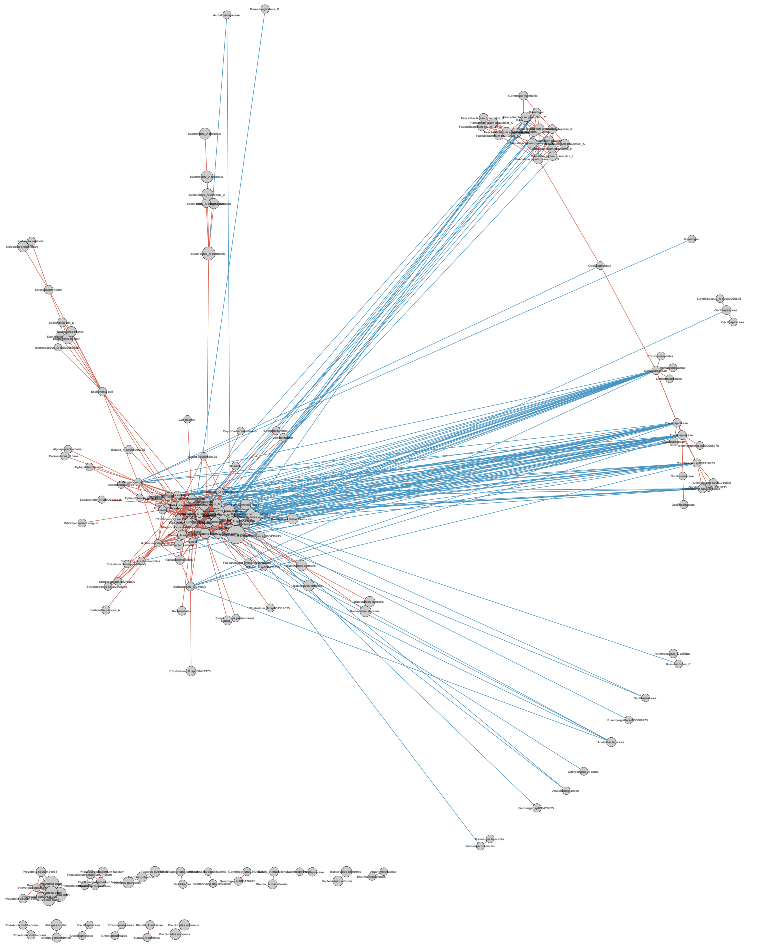

**Fig. S6 The co-abundance network of 184 genomes with unchanged correlations in $G_{M0}$, $G_{M3}$ and $G_{M15}$.** The correlations between the genomes were calculated using FastSpar. All significant correlations with P ≤ 0.001 were included. Node size indicates the average abundance of the genomes. Lines between nodes represent correlations, and red and blue colors indicate positive and negative correlations, respectively. Edge-weighted Spring Embedded Layout with correlation efficient as weight (-1 or 1) was applied to layout the network.

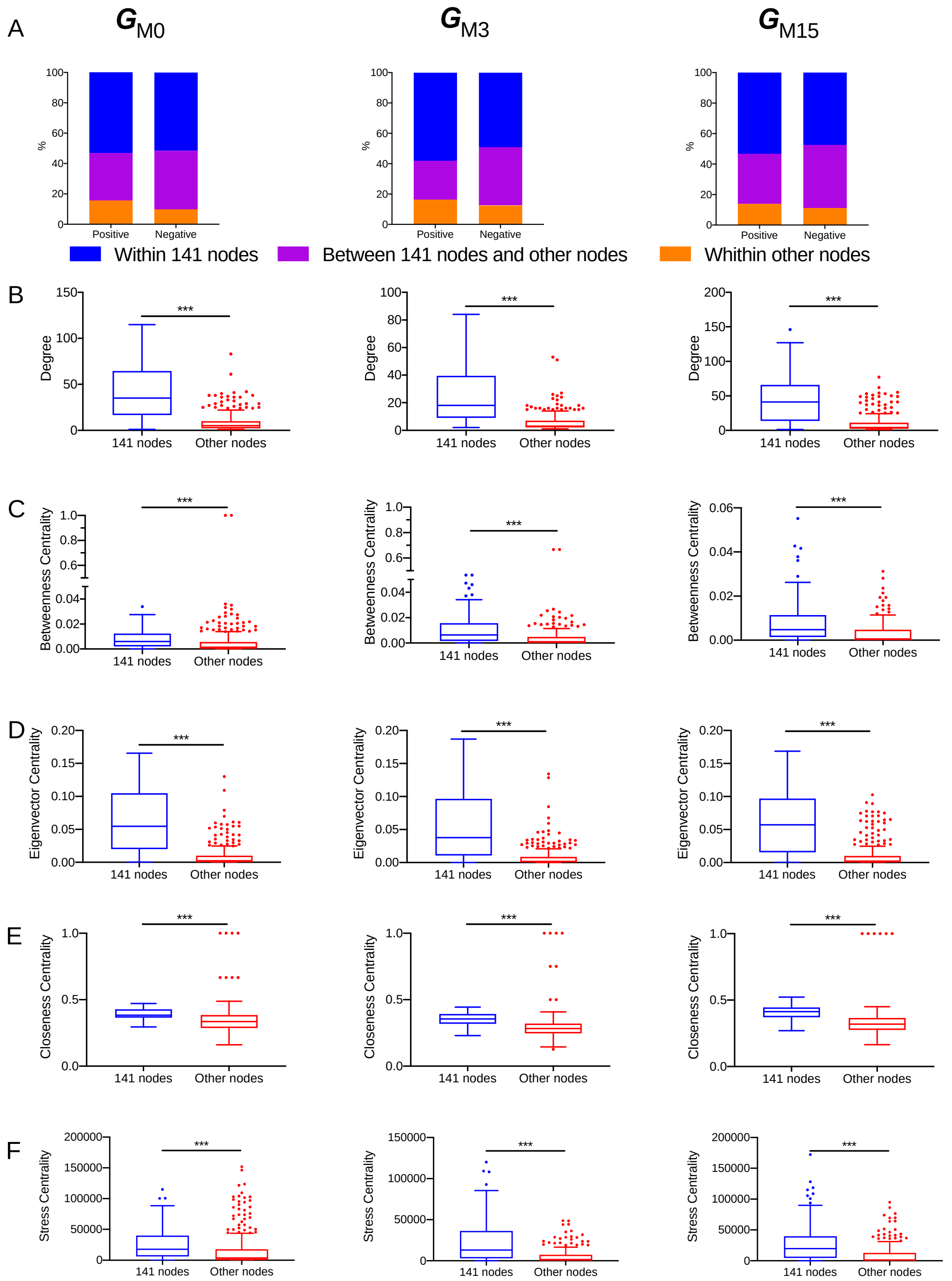


**Fig. S7 The 141 genomes contribute most of the interactions in the network.** (A) The stacked bar plot shows the distribution of positive and negative edges within the 141 genomes, between the 141 genomes and the other nodes, and within the other nodes. The 141 genomes had significantly higher degree (B), betweenness centrality (C), eigenvector centrality (D), closeness centrality (E) and stress centrality (F) than the rest of the nodes in the networks. Mann-Whitey test (two-sided) was performed. *** P < 0.0001

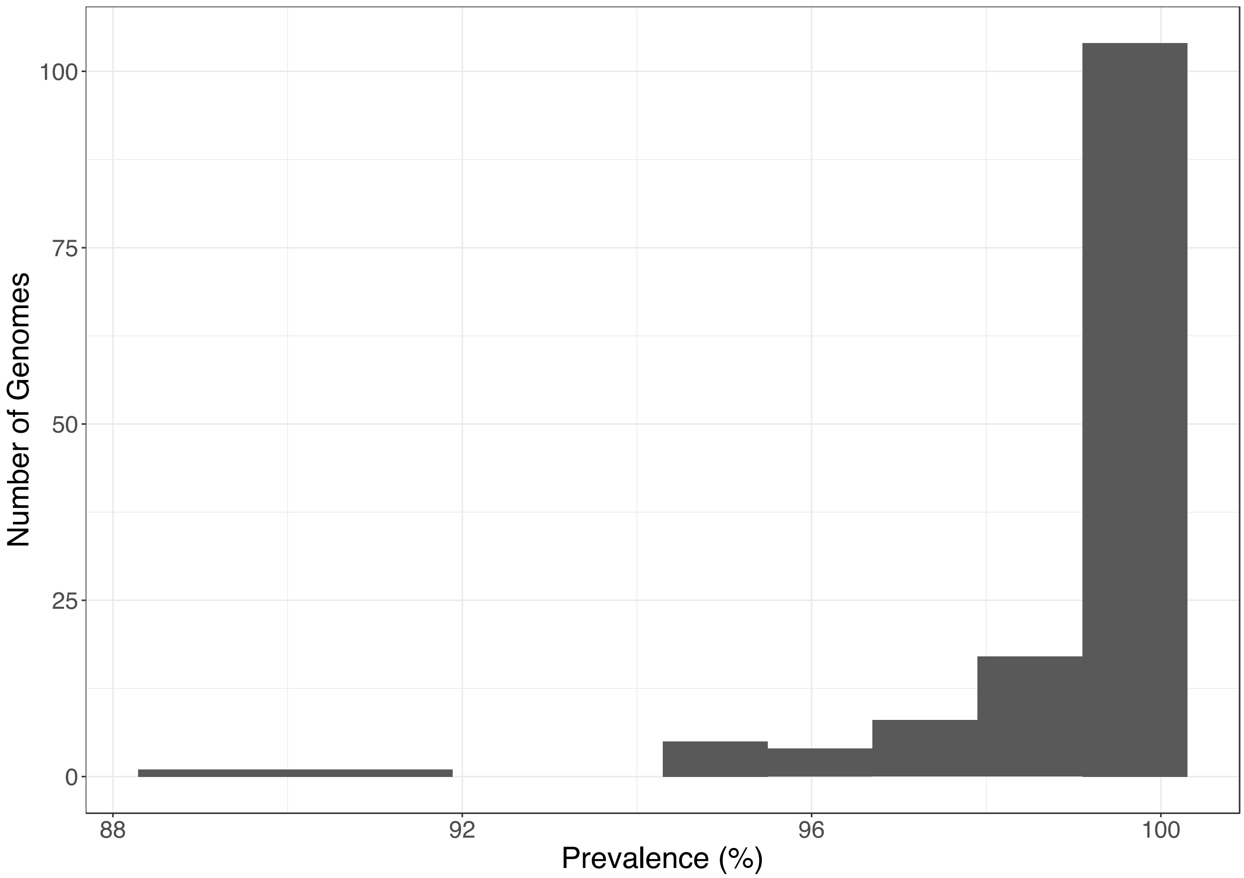


**Fig. S8 The 141 nodes were widely shared by the patients in the W group.** The histogram shows the distribution of genomes shared by the 74 patients with various prevalence.

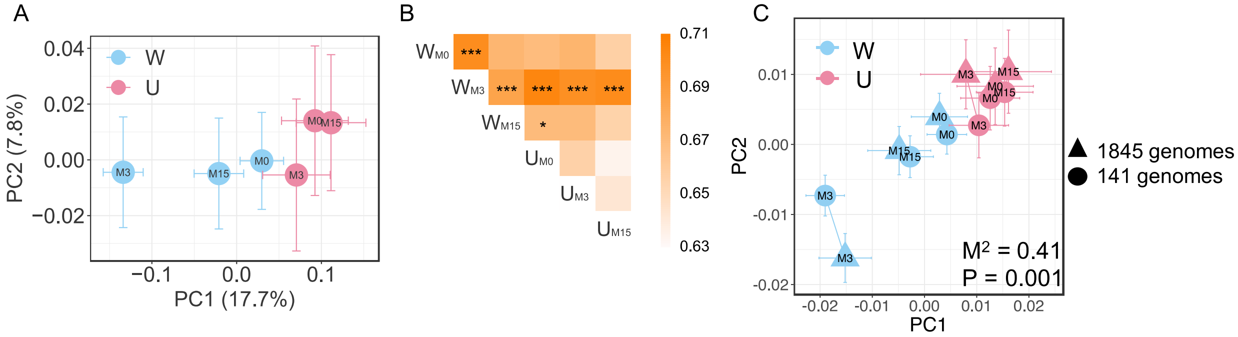


**Fig. S9 Similar beta-diversity pattern was found based on the 141 genomes as compared with that based on all the 1845 genomes.** (A) Global changes of the gut microbiome as shown by the principal coordinate analysis based on the Bray-Curtis distance with the abundance of the 141 genomes. (B) Average Bray-Curtis distance between the groups (B). PERMANOVA test (9,999 permutations) was performed to compare the groups. * $P < 0.05$ and *** $P < 0.001$. The color of the square showed the magnitude of average Bray-Curtis distance. (C) Procrustes analysis combing the principal coordinate analysis for 1845 genomes and 141 genomes based on Bray-Curtis distance.

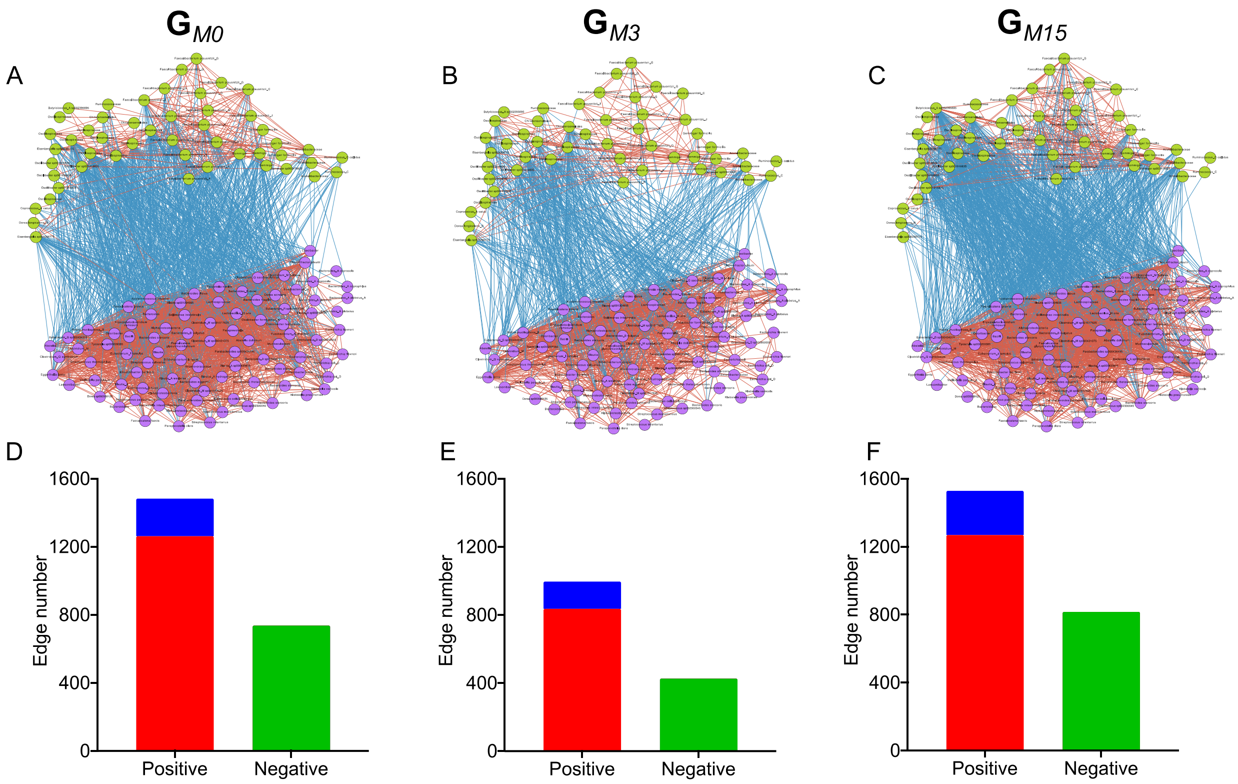


**Fig. S10 The 141 nodes organized themselves into two clusters with robust co-occurrence behavior within each cluster and can be recognized as potential ecological guilds.** (A-C) The co-abundance network within the 141 genomes. The correlations between the genomes were calculated using FastSpar, n = 67 patients. All significant correlations with P ≤ 0.001 were included. Edges between nodes represent correlations. Red and blue colors indicate positive and negative correlations, respectively. The color of the node represents the members in the two guilds: green for Guild 1 and purple for Guild 2. (D-F) The stacked bar plot shows the number of positive and negative edges within or between the guilds. Red, within Guild 1; Blue: within Guild 2; Green, between the two guilds.

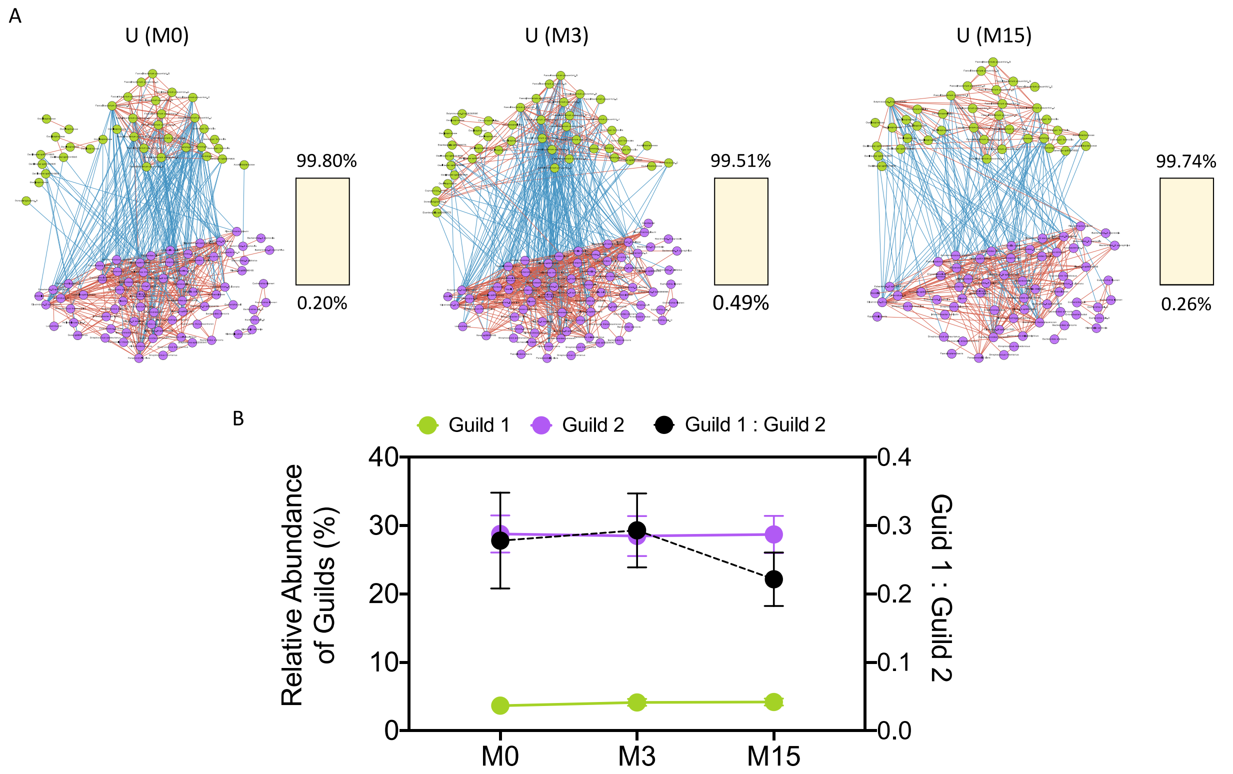


**Fig. S11 The seesaw networked genomes was detected in the U group and the balance between the guilds were not chagned.** (A)The correlations between the genomes were calculated using FastSpar at each timepoint of U group. All significant correlations with $P \leq 0.001$ were included. Lines between nodes represent correlations, and red and blue colors indicating positive and negative correlations, respectively. Node size indicates the average abundance of the genomes across the samples of patients with three timepoints data in U group n = 28. The color of the node represents the members in the two Guilds: green for Guild 1 and purple for Guild 2. The percentage of correlations followed the pattern in the seesaw network of the microbiome signature (i.e., positive edges within each guild, negative edges between the 2 guilds) was in yellow, and the ratio of correlations that were negative within each guild and positive between the guilds was in black of the 100% stacked bar. (B) Change of the total abundance of Guild 1, Guild 2, and their ratio across the trial in the U group. Friedman test followed by Nemenyi test was used to analyze the difference between time points. P values were > 0.05.

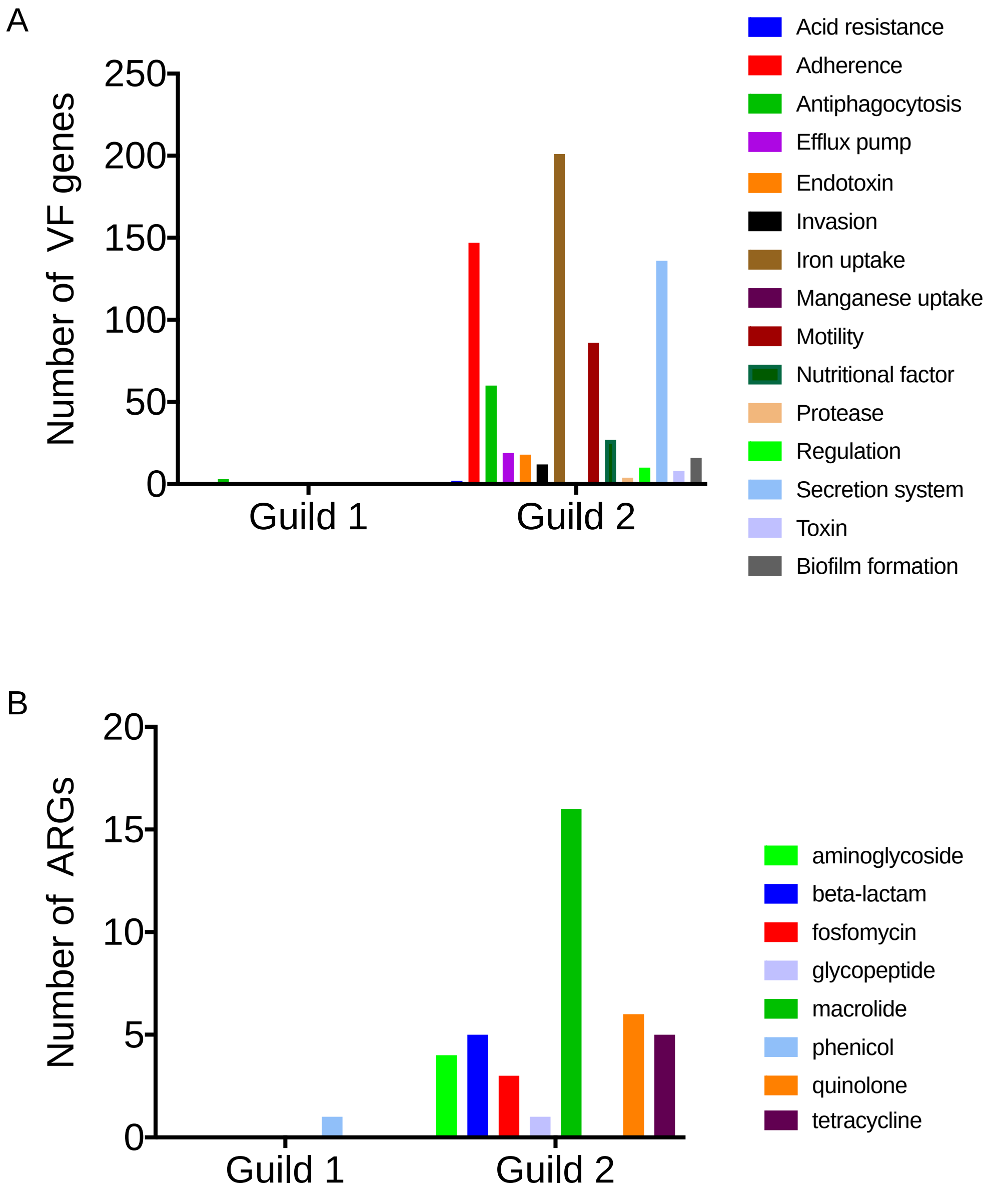


**Fig. S12. Genomes in Guild 1 had much lower genetic capacity for pathogenicity and antibiotic resistance.** (A) The bar plot shows the number of genes encoding virulence factors (VF) and classes of VFs. (B) The bar plot shows the number of ARGs and the corresponding antibiotic resistance types.

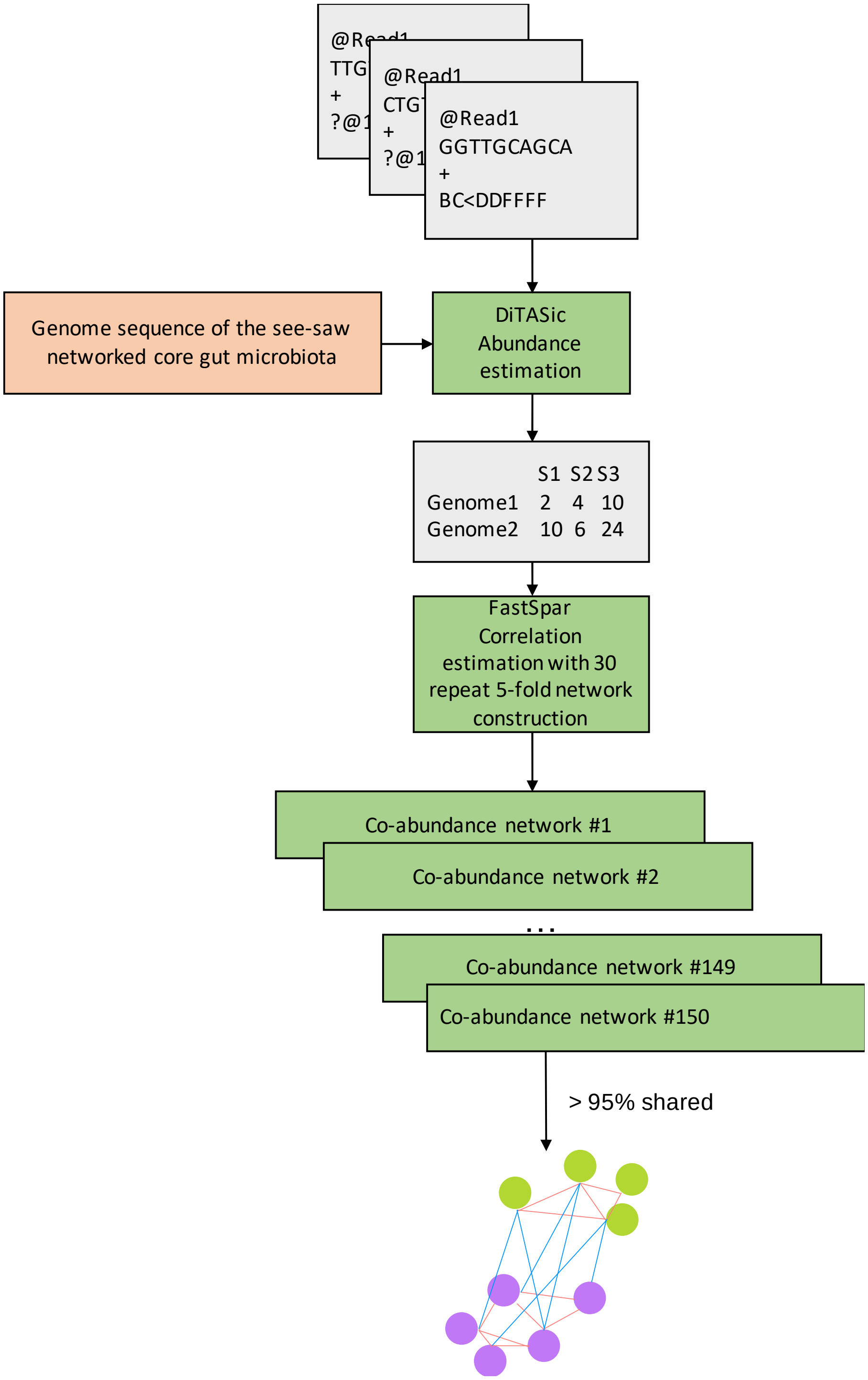
@Read1
TTG
+
?@1
@Read1
CTG
+
?@1
@Read1
GGTTGCAGCA
+
BC<DDFFFF
Genome sequence of the see-saw networked core gut microbiota
DiTASic Abundance estimation
S1 S2 S3
Genome1 2 4 10
Genome2 10 6 24
FastSpar Correlation estimation with 30 repeat 5-fold network construction
Co-abundance network #1
Co-abundance network #2
...
Co-abundance network #149
Co-abundance network #150
> 95% shared

**Fig. S13. The workflow for validating the microbiome signature in other datasets.**

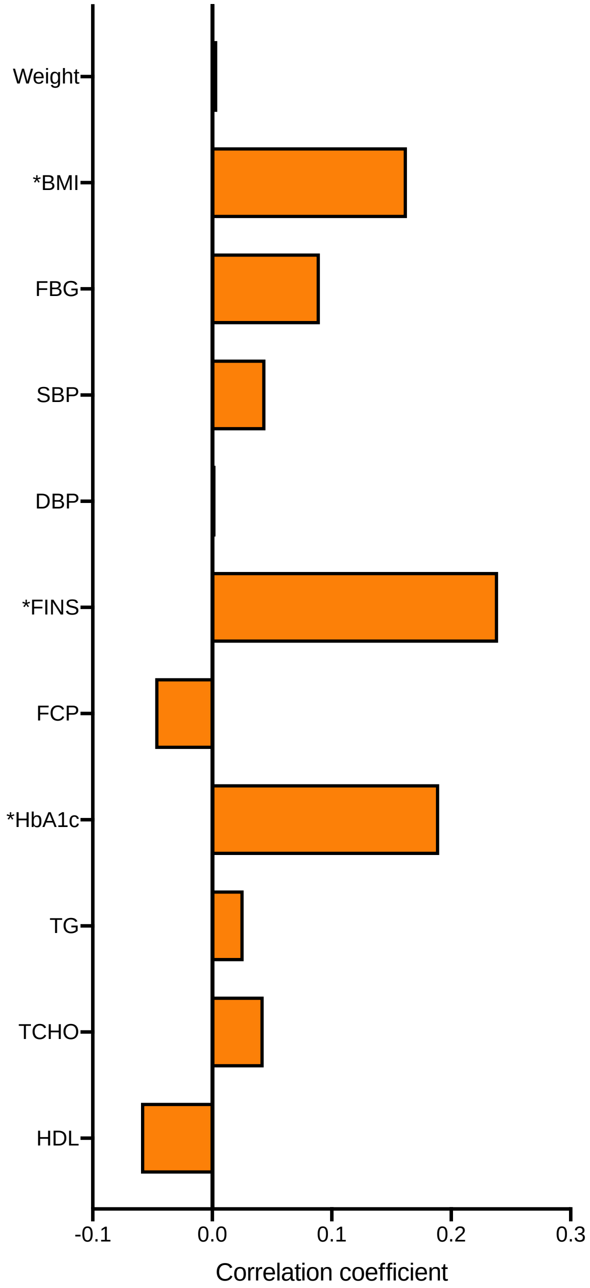


**Fig. S14. The microbiome signature is associated with host phenotype in an independent type 2 diabetes dataset (Qin 2012, et al.).** Random Forest regression with leave-one-out cross-validation was used to explore the associations between the microbiome signature and clinical parameters. The bar plot shows the Pearson's correlations coefficient between the predicted values from the regression model and measured values. The asterisk before the parameter's name shows the significance of the Pearson's correlations. P values were adjusted by Benjamini & Hochberg's method.

* adjusted P < 0.05. BMI: body mass index, FBG: fasting blood glucose, SBP: systolic blood glucose, DBP: diastolic blood pressure, FINS: fasting serum insulin, FCP: fasting serum C-peptide, TG: triglyceride, TCHO: total cholesterol, HDL: High density lipoprotein. N = 272.

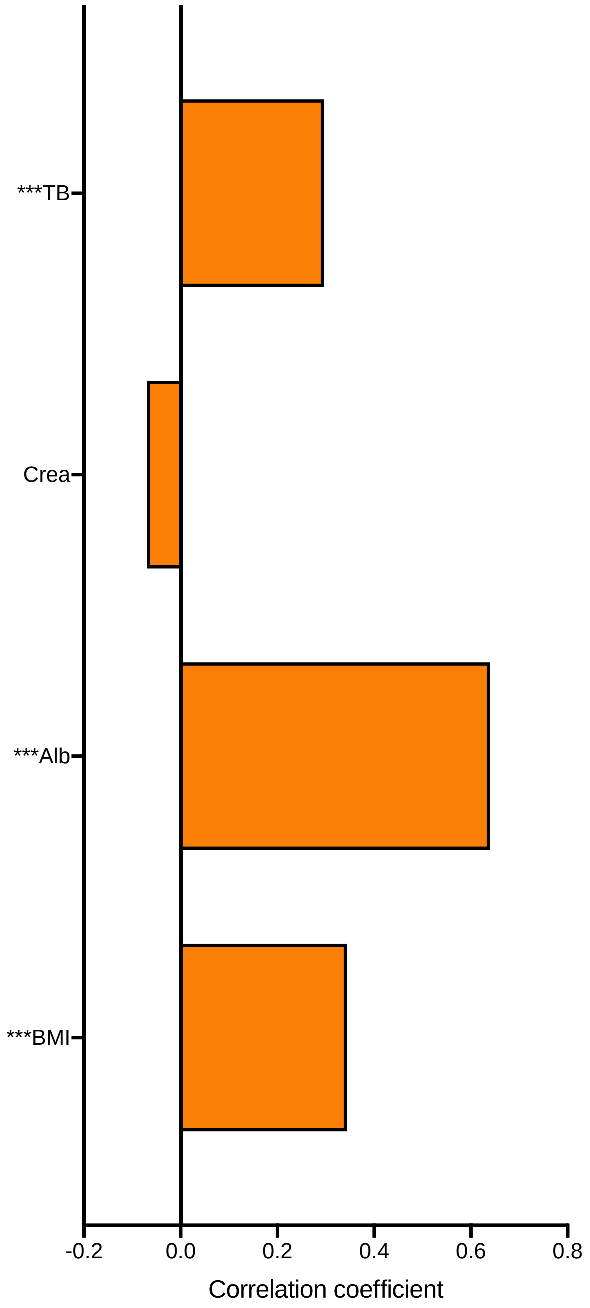

**Fig. S15. The microbiome signature is associated with host phenotypes in a liver cirrhosis dataset (Qin 2014, et al.)**. Random Forest regression with leave-one-out cross-validation was used to explore the associations between the microbiome signature and the clinical parameters. The bar plot shows the Pearson's correlations coefficient between the predicted values from the regression model and measured values. The asterisk before the parameter's name shows the significance of the Pearson's correlations. P values were adjusted by Benjamini & Hochberg's method. *** adjusted P < 0.001. TB: total bilirubin, Crea: creatinine level, Alb: albumin level, BMI: Body mass index. N = 167.

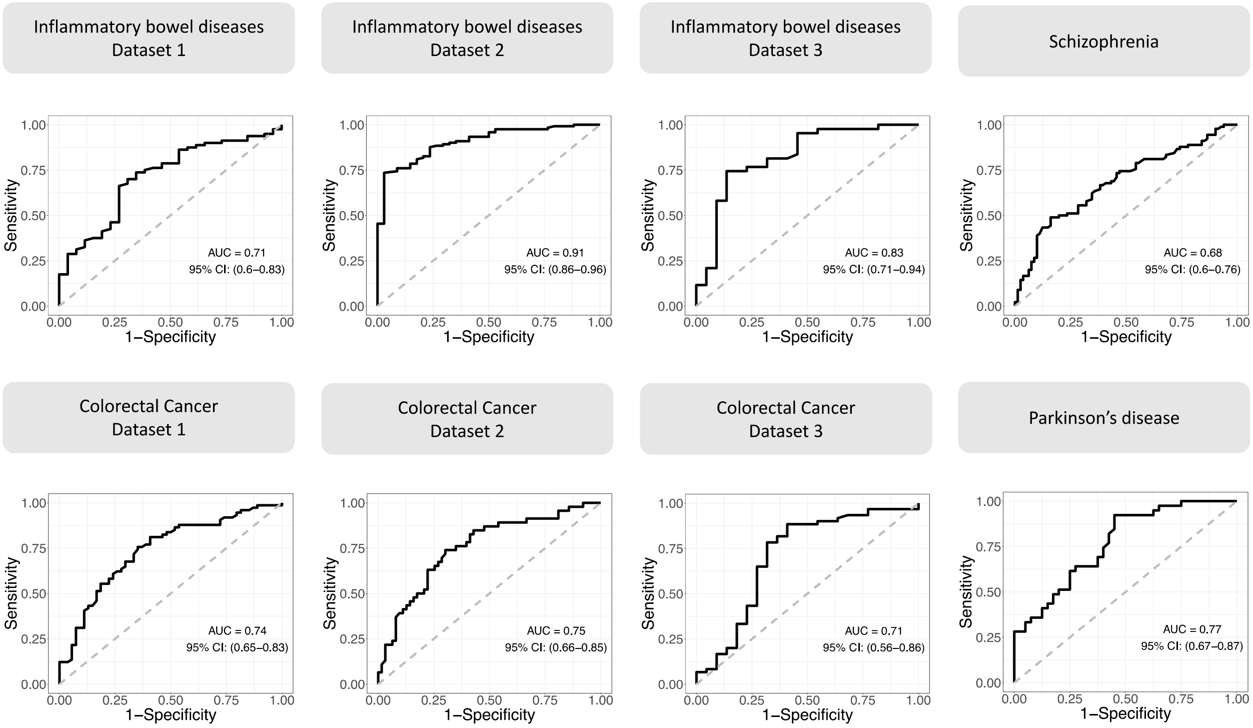


**Fig. S16. The discriminative power of microbiome signature as biomarkers to classify healthy subjects vs. patients in datasets on more diseases across ethnicity and geography.** The microbiome signature supports predictive classification models for 8 other independent datasets. The area under the ROC curve (AUC) of the Random Forest classifier based on the 141 genomes in the microbiome signature to classify control and patients in each dataset. Leave-one-out cross validation was applied. Inflammatory bowel disease (IBD) Dataset 1[1]: Control n = 26, IBD n = 80; IBD Dataset 2[2]: Control n = 34, IBD n = 121; IBD Dataset 3[2]: Control n = 22, IBD n =43. Colorectal Cancer (CRC) Dataset 1[3]: Control n = 54, CRC n = 74; CRC Dataset 2[4]: Control n = 63, CRC n = 46; CRD Dataset 3[5]: Control n = 60, CRC n = 22; Schizophrenia[6]: Control = 81, Schizophrenia n = 90; Parkinson's Disease[7]: Control n = 40, Parkinson's Disease n = 39. The sample ids were listed in Table S8.

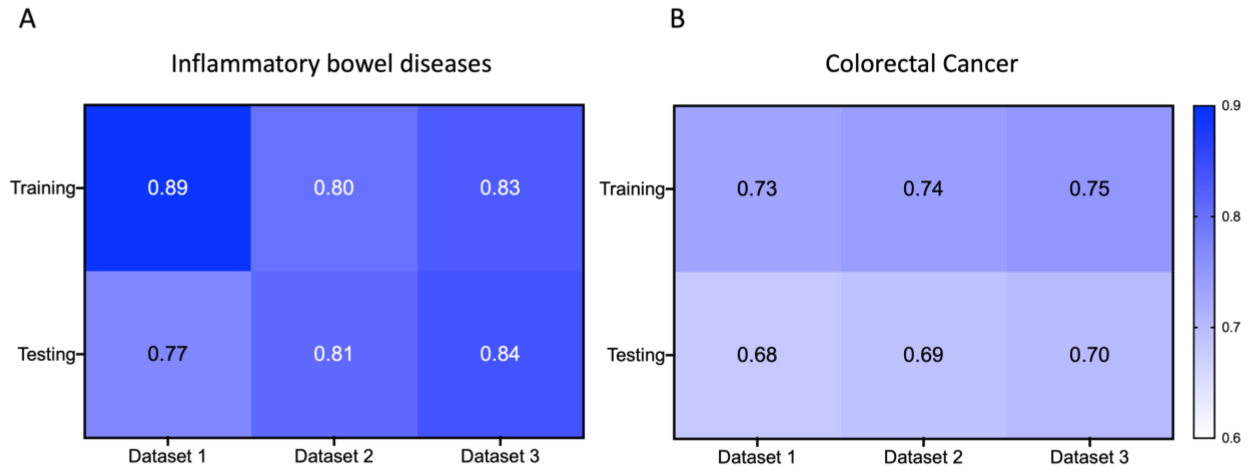


**Fig. S17 The discriminative power of microbiome signature as biomarkers to classify healthy subjects vs. patients in datasets of inflammatory bowel diseases (IBD) and colorectal cancer (CRC) with leave-one-cohort-out (LOCO) analysis.** LOCO analysis was performed on IBD (A) and CRC (B), which both had multiple datasets included in our study. In IBD/CRC dataset, the abundances of the genomes were adjusted by MMUPHin to correct batch effect. In LOCO analysis, one dataset was kept for testing and the other two were used for model training using Random Forest classifier based on the 141 genomes in the microbiome with leave-one-out cross-validation. The model was then applied to the testing dataset. The heatmaps show the area under the ROC curve (AUC) both in training and testing results. Inflammatory bowel disease (IBD) Dataset 1[1]: Control n = 26, IBD n = 80; IBD Dataset 2[2]: Control n = 34, IBD n = 121; IBD Dataset 3[2]: Control n = 22, IBD n =43. Colorectal Cancer (CRC) Dataset 1[3]: Control n = 54, CRC n = 74; CRC Dataset 2[4]: Control n = 63, CRC n = 46; CRD Dataset 3[5]: Control n = 60, CRC n = 22;

### Table S1 Antidiabetic medication.

[In the Excel file]

### Table S2 Raw and high-quality reads of each sample.

[In the Excel file]

### Table S3 Clinical parameters during intervention in the W and U group.

| | W | | | U | | | P value (W vs U) | | |
|---|---|---|---|---|---|---|---|---|---|
| | M0 | M3 | M15 | M0 | M3 | M15 | M0 | M3 | M15 |
| **General Information** | | | | | | | | | |
| Age (years) | 60.91 ± 0.83 (74) | / | / | 58.81 ± 1.35 (36) | / | / | 0.1865 | / | / |
| Duration of diabetes (years) | 9.83 ± 0.8 (72) | / | / | 10.94 ± 1.03 (35) | / | / | 0.2853 | / | / |
| Gender (Males/Females) | 23/51 | 23/51 | 22/45 | 14/22 | 14/22 | 11/17 | 0.5497 | / | / |
| **Glycometabolism** | | | | | | | | | |
| HbA1c (%) | 8.17 ± 0.14$^{a}$ (74) | 6.85 ± 0.08$^{c}$ (74) | 7.74 ± 0.13$^{b}$ (67) | 8.08 ± 0.2 (36) | 7.64 ± 0.16 (36) | 7.67 ± 0.18 (28) | 0.7163 | 0.0001 | 0.7070 |
| FBG (mmol/L) | 9.57 ± 0.26$^{a}$ (74) | 7.18 ± 0.14$^{c}$ (74) | 8.28 ± 0.19$^{b}$ (67) | 8.66 ± 0.23$^{a}$ (36) | 7.68 ± 0.36$^{b}$ (36) | 7.84 ± 0.33$^{ab}$ (28) | 0.0524 | 0.5429 | 0.4700 |
| MTT Glucose AUC (mmol/L/min) | 2800.65 ± 64.48$^{a}$ (72) | 2252.5 ± 51.15$^{c}$ (74) | 2412.55 ± 64.15$^{b}$ (67) | 2478.81 ± 67.43 (36) | 2400.67 ± 77.91 (36) | 2368.43 ± 74.63 (28) | 0.0078 | 0.1938 | 0.7012 |
| MTT C-Peptide AUC (ng/mL/min) | 874.73 ± 43.41 (72) | 790.15 ± 36.93 (74) | 814.95 ± 42 (67) | 937.85 ± 71.43$^{a}$ (36) | 789.13 ± 55.91$^{b}$ (36) | 800.99 ± 65.01$^{b}$ (28) | 0.4171 | 0.9670 | 0.8575 |
| HOMA-IR | 4.12 ± 0.24$^{a}$ (74) | 2.93 ± 0.07$^{c}$ (74) | 3.41 ± 0.1$^{b}$ (67) | 4.06 ± 0.2$^{a}$ (36) | 3.43 ± 0.17$^{b}$ (36) | 3.61 ± 0.21$^{ab}$ (28) | 0.5220 | 0.0192 | 0.5650 |
| HOMA-β | 37.35 ± 2.5$^{a}$ (74) | 45.17 ± 2.76$^{b}$ (74) | 39.78 ± 2.36$^{b}$ (67) | 45.75 ± 4.09$^{a}$ (36) | 60.61 ± 8.66$^{ab}$ (36) | 139.87 ± 90.46$^{b}$ (28) | 0.0809 | 0.3640 | 0.0657 |
| **Anthropometric markers** | | | | | | | | | |
| BMI (kg/m$^2$) | 25.81 ± 0.39$^{a}$ (74) | 24.55 ± 0.36$^{b}$ (74) | 25.98 ± 0.44$^{a}$ (66) | 26.82 ± 0.51$^{a}$ (36) | 26.27 ± 0.5$^{b}$ (36) | 27.43 ± 0.64$^{a}$ (28) | 0.0670 | 0.0017 | 0.0372 |
| BW (Kg) | 66.58 ± 1.34$^{a}$ (74) | 63.32 ± 1.26$^{b}$ (74) | 66.91 ± 1.41$^{a}$ (66) | 71.25 ± 1.71$^{a}$ (36) | 69.82 ± 1.72$^{b}$ (36) | 72.25 ± 2.16$^{a}$ (28) | 0.0176 | 0.0013 | 0.0422 |
| SBP (mmHg) | 132.39 ± 1.48$^{a}$ (74) | 122.66 ± 2.07$^{b}$ (73) | 127.52 ± 2.03$^{a}$ (66) | 131.06 ± 2.82 (33) | 128.38 ± 2.39 (32) | 125.35 ± 2.81 (26) | 0.3245 | 0.0342 | 0.7318 |
| DBP (mmHg) | 73.78 ± 1.31$^{a}$ (74) | 71.21 ± 1.43$^{ab}$ (73) | 69.55 ± 1.45$^{b}$ (66) | 76.97 ± 2.27 (33) | 74.12 ± 2.13 (32) | 71.77 ± 2.29 (26) | 0.5615 | 0.2681 | 0.5876 |
| WC (cm) | 86.73 ± 1.13$^{a}$ (74) | 84.43 ± 1.06$^{b}$ (74) | 87.05 ± 1.17$^{a}$ (65) | 92.83 ± 1.42 (35) | 90.97 ± 1.43 (36) | 93.75 ± 1.78 (28) | 0.0006 | 0.0006 | 0.0031 |
| HC (cm) | 97.36 ± 0.8$^{a}$ (74) | 93.35 ± 0.85$^{c}$ (74) | 95.23 ± 0.82$^{b}$ (65) | 97.89 ± 0.78 (35) | 98.28 ± 0.94 (36) | 97.64 ± 1.09 (28) | 0.3961 | 0.0004 | 0.0766 |
| WHR | 0.89 ± 0.01$^{a}$ (74) | 0.9 ± 0.01$^{a}$ (74) | 0.91 ± 0.01$^{b}$ (65) | 0.95 ± 0.01 (35) | 0.92 ± 0.01 (36) | 0.96 ± 0.01 (28) | 0.0001 | 0.0469 | 0.0019 |
| **Inflammatory markers** | | | | | | | | | |
| TNF-α (pg/ml) | 1.28 ± 0.05 (74) | 1.29 ± 0.06 (74) | 1.75 ± 0.2 (67) | 1.26 ± 0.06$^{a}$ (36) | 1.28 ± 0.08$^{a}$ (36) | 3.52 ± 0.65$^{b}$ (28) | 0.7477 | 0.7915 | 0.0010 |
| WBC (10$^9$/L) | 5.88 ± 0.17$^{a}$ (74) | 5.5 ± 0.15$^{b}$ (74) | 6.17 ± 0.21$^{a}$ (67) | 5.76 ± 0.26$^{a}$ (36) | 6.14 ± 0.28$^{ab}$ (36) | 6.12 ± 0.33$^{b}$ (28) | 0.6672 | 0.0544 | 0.8639 |
| CRP (mg/L) | 1.56 ± 0.22$^{a}$ (73) | 1.14 ± 0.17$^{b}$ (74) | 0.77 ± 0.15$^{b}$ (65) | 1.21 ± 0.25 (36) | 1.29 ± 0.19 (35) | 0.84 ± 0.11 (28) | 0.1796 | 0.2337 | 0.0171 |
| LBP (ug/ml) | 5.5 ± 0.17$^{a}$ (74) | 5.75 ± 0.1$^{c}$ (74) | 4.93 ± 0.27$^{b}$ (67) | 5.68 ± 0.21$^{a}$ (36) | 5.55 ± 0.22$^{ab}$ (36) | 4.93 ± 0.22$^{b}$ (28) | 0.2609 | 0.0804 | 0.2254 |
| **Adipocytokines** | | | | | | | | | |
| Adiponectin (ng/ml) | 13.96 ± 1.48$^{a}$ (64) | 13.76 ± 1.45$^{a}$ (64) | 12.2 ± 1.46$^{b}$ (64) | 15.02 ± 2.42 (26) | 13.37 ± 2.29 (26) | 12.48 ± 2.18 (26) | 0.7520 | 0.6724 | 0.7792 |
| Leptin (pg/ml) | 876.53 ± 83.62$^{a}$ (64) | 646.74 ± 67.17$^{b}$ (64) | 878.69 ± 99.86$^{a}$ (64) | 1231.46 ± 170.09 (26) | 1230.33 ± 194.76 (26) | 1540.24 ± 228.52 (26) | 0.0551 | 0.0006 | 0.0013 |
| **Lipid metabolism** | | | | | | | | | |
| TC (mmol/L) | 5.18 ± 0.13$^{a}$ (74) | 4.24 ± 0.1$^{b}$ (74) | 4.59 ± 0.11$^{b}$ (67) | 4.82 ± 0.17$^{a}$ (36) | 4.44 ± 0.14$^{b}$ (36) | 4.54 ± 0.17$^{ab}$ (28) | 0.0755 | 0.2903 | 0.8002 |
| TG (mmol/L) | 1.27 ± 0.06$^{a}$ (69) | 1.04 ± 0.06$^{b}$ (72) | 1.22 ± 0.07$^{a}$ (64) | 1.48 ± 0.09 (33) | 1.45 ± 0.09 (32) | 1.38 ± 0.11 (28) | 0.0539 | 0.0001 | 0.1653 |
| Lpa (mg/L) | 235.58 ± 26.79$^{a}$ (74) | 201.86 ± 20.02$^{a}$ (74) | 163.72 ± 24.56$^{b}$ (67) | 178.19 ± 26.9$^{a}$ (36) | 165.44 ± 24.7$^{a}$ (36) | 110.89 ± 16.44$^{b}$ (28) | 0.2298 | 0.1091 | 0.4625 |
| HDL (mmol/L) | 1.37 ± 0.04$^{a}$ (74) | 1.31 ± 0.04$^{a}$ (74) | 1.48 ± 0.04$^{b}$ (67) | 1.23 ± 0.04$^{a}$ (36) | 1.17 ± 0.04$^{c}$ (36) | 1.34 ± 0.06$^{b}$ (28) | 0.0318 | 0.0218 | 0.1484 |
| APOA (g/L) | 1.53 ± 0.03$^{a}$ (74) | 1.59 ± 0.04$^{a}$ (74) | 1.43 ± 0.03$^{b}$ (67) | 1.42 ± 0.05 (36) | 1.47 ± 0.06 (36) | 1.35 ± 0.06 (28) | 0.0729 | 0.1193 | 0.2071 |
| LDL (mmol/L) | 2.69 ± 0.09$^{ab}$ (74) | 2.52 ± 0.08$^{b}$ (74) | 2.84 ± 0.09$^{a}$ (67) | 2.58 ± 0.11$^{a}$ (36) | 2.68 ± 0.11$^{a}$ (36) | 2.96 ± 0.13$^{b}$ (28) | 0.4934 | 0.2732 | 0.5032 |
| APOB (g/L) | 1 ± 0.03$^{a}$ (74) | 0.79 ± 0.02$^{b}$ (74) | 0.98 ± 0.03$^{a}$ (67) | 0.98 ± 0.04$^{a}$ (36) | 0.87 ± 0.03$^{b}$ (36) | 1.01 ± 0.04$^{a}$ (28) | 0.5407 | 0.0749 | 0.5403 |
| **Kidney function** | | | | | | | | | |

| | | | | | | | | | |
|---|---|---|---|---|---|---|---|---|---|
| GFR (mL/min per 1.73 $m^2$) | 120.74 ± 2.81$^{a}$ (74) | 119.62 ± 2.19$^{ab}$ (74) | 129.43 ± 3.2$^{b}$ (67) | 130.01 ± 4.37$^{a}$ (36) | 125.15 ± 4.16$^{a}$ (36) | 115.27 ± 5$^{b}$ (28) | 0.0166 | 0.1611 | 0.0166 |
| CysC (mg/L) | 0.65 ± 0.01$^{ab}$ (74) | 0.67 ± 0.01$^{b}$ (74) | 0.62 ± 0.01$^{a}$ (67) | 0.65 ± 0.03$^{a}$ (36) | 0.68 ± 0.03$^{b}$ (36) | 0.68 ± 0.04$^{ab}$ (28) | 0.4951 | 0.8935 | 0.3229 |
| ACR (ug/mg) | 28.27 ± 4.78 (68) | 21.85 ± 3.97 (69) | 24.3 ± 4.3 (63) | 34.45 ± 8.28 (33) | 39.67 ± 9.5 (35) | 34.41 ± 8.72 (25) | 0.3929 | 0.0330 | 0.0996 |
| **Atherosclerosis** | | | | | | | | | |
| IMT (mm) | 0.88 ± 0.01$^{a}$ (73) | 0.83 ± 0.01$^{b}$ (74) | 0.88 ± 0.02$^{a}$ (67) | 0.78 ± 0.02 (36) | 0.83 ± 0.02 (35) | 0.89 ± 0.03 (28) | 0.0034 | 0.9715 | 0.7315 |
| **Diabetic autonomic neuropathy** | | | | | | | | | |
| DAN score | 2.47 ± 0.12$^{a}$ (74) | 2 ± 0.11$^{ab}$ (73) | 1.87 ± 0.13$^{b}$ (66) | 1.89 ± 0.17 (33) | 1.92 ± 0.15 (32) | 1.94 ± 0.18 (26) | 0.0150 | 0.7430 | 0.8609 |
| MHR | 75.76 ± 1$^{a}$ (74) | 72.2 ± 0.87$^{b}$ (74) | 72.55 ± 0.88$^{b}$ (67) | 76.5 ± 1.48 (36) | 74.11 ± 1.61 (36) | 75.57 ± 2.06 (28) | 0.7692 | 0.3387 | 0.1057 |
| SDNN (ms) | 105.19 ± 2.75$^{a}$ (74) | 112.54 ± 2.67$^{b}$ (74) | 115.75 ± 3.29$^{b}$ (67) | 120.58 ± 5.95 (36) | 123.81 ± 5.34 (36) | 123.68 ± 6.39 (28) | 0.0245 | 0.0651 | 0.3074 |
| SDANN (ms) | 97.86 ± 2.76 (74) | 103.64 ± 2.66 (74) | 106.49 ± 3.68 (55) | 112.33 ± 5.6 (36) | 116.64 ± 5.36 (36) | 117.12 ± 6.68 (24) | 0.0320 | 0.0372 | 0.2142 |
| SDNNIndex (ms) | 41.22 ± 1.28$^{a}$ (74) | 42.97 ± 1.12$^{b}$ (74) | 41.93 ± 1.2$^{ab}$ (67) | 41.64 ± 1.71 (36) | 41.11 ± 1.96 (36) | 40.21 ± 2.41 (28) | 0.4595 | 0.7258 | 0.4163 |
| rMSSD (ms) | 23.88 ± 1.15 (74) | 24.72 ± 1.13 (74) | 25.22 ± 1.44 (67) | 22.44 ± 1.23 (36) | 20.94 ± 1.2 (36) | 21.39 ± 1.73 (28) | 0.7042 | 0.0537 | 0.0553 |
| pNN50 (%) | 4.12 ± 0.64 (74) | 4.74 ± 0.6 (74) | 5.27 ± 0.72 (67) | 4.06 ± 0.7 (36) | 3.33 ± 0.63 (36) | 4 ± 1.08 (28) | 0.7453 | 0.1112 | 0.1115 |
| TP ($ms^2$) | 1656.32 ± 88.06$^{ab}$ (74) | 1887.86 ± 104.32$^{b}$ (74) | 1688.18 ± 97.78$^{a}$ (59) | 1814.81 ± 131.57 (36) | 1859.14 ± 161.38 (36) | 1664.77 ± 193.33 (28) | 0.2663 | 0.9011 | 0.4870 |
| VLF ($ms^2$) | 1253.8 ± 67.77$^{ab}$ (74) | 1431.83 ± 82.41$^{b}$ (74) | 1231.65 ± 70.21$^{a}$ (59) | 1392.23 ± 102.38 (36) | 1446.21 ± 127.71 (36) | 1280.57 ± 152.58 (28) | 0.2582 | 0.6442 | 0.6334 |
| LF ($ms^2$) | 262.37 ± 16.37$^{ab}$ (74) | 292.98 ± 19.73$^{b}$ (74) | 267.34 ± 20.71$^{a}$ (59) | 287.74 ± 26.17 (36) | 289.24 ± 30.09 (36) | 247.05 ± 31.31 (28) | 0.4094 | 0.8659 | 0.3856 |
| HF ($ms^2$) | 126.3 ± 10.21 (74) | 144.71 ± 11.32 (74) | 161.53 ± 21.46 (59) | 121.37 ± 14.43 (36) | 114.01 ± 12.89 (36) | 119.97 ± 23.25 (28) | 0.8186 | 0.1239 | 0.0466 |
| **Diabetic peripheral neuropathy** | | | | | | | | | |
| DPN score | 0.96 ± 0.09 (74) | 0.77 ± 0.1 (74) | 0.93 ± 0.1 (67) | 1.03 ± 0.14 (36) | 0.92 ± 0.13 (36) | 1.39 ± 0.16 (28) | 0.6851 | 0.3515 | 0.0148 |
| Michigan score | 1.76 ± 0.21 (74) | 1.46 ± 0.16 (74) | 1.87 ± 0.22 (67) | 2.22 ± 0.35 (36) | 1.79 ± 0.29 (36) | 2.45 ± 0.42 (28) | 0.1941 | 0.3844 | 0.3001 |

The data are showed as mean ± S.E.M (N). Friedman test followed by Nemenyi post-hoc test was used for intra-group comparisons, means with the same letter (a, b, or c) are not significantly different, with different letters are significantly different ($P < 0.05$). Mann-Whitney test (two-sided) was used for comparisons between W and U at the same time point. FBG, fasting blood glucose; MTT Glucose AUC, area under the curve (AUC) of glucose in meal tolerance test; MTT C-Peptide AUC , area under the curve (AUC) of C-Peptide in meal tolerance test; HOMA-IR = 1.5 + FBG * Fasting-C-Peptide / 2800; HOMA-β = 0.27 * Fasting-C-Peptide / (FBG - 3.5); BMI, body mass index; BD, body weight; SBP, systolic blood pressure; DBP, diastolic blood pressure; WC, waist circumference; HP, hip circumference; WHR, waist to hip ratio; TNF-α, tumor necrosis factor-α; WBC, white blood cell count; CRP, C-reactive protein; LBP, lipopolysaccharide-binding protein; TC, total cholesterol; TG, triglyceride; Lpa, lipoprotein a; HDL, high-density lipoprotein; APOA, apolipoprotein A; LDL, low-density lipoprotein; APOB, apolipoprotein B; GFR, glomerular filtration rate; CysC, Cystatin C; ACR, urinary microalbumin to creatinine ratio; IMT, intima-media thickness; DAN, diabetic autonomic neuropathy score; MHR, mean heart rate; SDNN, standard deviation of NN intervals; SDANN, standard deviation of the average NN intervals calculated over 5 minutes; SDNNIndex, mean of standard deviation of NN intervals for 5-minute segments; rMSSD, root-mean-square of the differences of successive NN intervals; pNN50, percentage of the interval differences of successive NN intervals greater than 50 ms; TP, total power; VLF, very low frequency power; LF, low frequency power; HF, high frequency power; DPN, diabetic peripheral neuropathy score.

**Table S4 Topological parameters of each network in the W group.**

| | $G_{M0}$ | $G_{M3}$ | $G_{M15}$ |
|---|---|---|---|
| Order (***S***, total number of nodes) | 442 | 421 | 429 |
| Size (***L***, total number of edges) | 4231 | 2587 | 4592 |
| # of negative correlations | 1432 | 869 | 1722 |
| # of positive correlations | 2799 | 1718 | 2870 |
| Connectance | 0.04341224 | 0.0292614 | 0.05001852 |
| Avg. number of neighbors | 19.1447964 | 12.2897862 | 21.40792541 |
| Clustering coefficient | 0.35579511 | 0.27844768 | 0.379733602 |
| Network diameter | 9 | 11 | 9 |
| Network centralization | 0.21834673 | 0.17155359 | 0.292466458 |
| Shortest paths | 187936 | 170172 | 178512 |
| Characteristic path length | 3.04453644 | 3.42079778 | 3.004795196 |
| Network heterogeneity | 1.25224147 | 1.31946401 | 1.294432497 |

**Table S5 The table corresponds to Figure 3A.**
[In the Excel file]

**Table S6 The genome quality assessed by CheckM.**
[In the Excel file]

**Table S7 The taxonomic assignment of genomes by GT-DBTK.**
[In the Excel file]

**Table S8 The information of samples for validation in independent datasets.**
[In the Excel file]
